# Synergistic action of different molecular mechanisms causes striking levels of insecticide resistance in the malaria vector *Anopheles gambiae*

**DOI:** 10.64898/2026.03.23.713604

**Authors:** Mengling Chen, Latifa Remadi, Dimitra Tsakireli, Emmanouil Kokkas, Sofia Balaska, Sibora Teta, Jocelyn M. F. Ooi, Janet Hemingway, Mark J.I. Paine, Gareth J. Lycett, John Vontas, Linda Grigoraki

## Abstract

Intensifying insecticide resistance in the malaria vector *Anopheles gambiae* poses a serious threat to the hard-won gains in reducing malaria deaths in Africa. The genetic basis of insecticide resistance is often complex, involving multiple genes and mutations. However, we still lack a clear understanding of how each mechanism contributes to overall resistance and how highly resistant phenotypes arise. In this study we generated a suite of transgenic *An. gambiae* strains carrying either individual mechanisms or combinations that frequently co-occur in nature. We show that co-overexpression of different classes of detoxification enzymes (CYP6P3, CYP6M2, CYP9K1, ABCH2, GSTE2 and COEAE6G), as well as the overexpression of detoxification enzymes in the presence of target site resistance mutations, can lead to substantially greater levels of resistance. Our findings suggest that increased resistance strength is a primary driver for selection of multi-mechanism resistance and are transformative for the scientific insight required to design robust molecular diagnostics for timely and reliable resistance detection in the field. We further show that P450 based resistance can constitute an Achilles heel for highly resistant mosquitoes, making them more vulnerable to pro-insecticides; compounds that typically require P450 activation. Our results advance our understanding of the mechanistic basis of insecticide resistance and have important implications for the design and implementation of effective and evidence-based resistance management strategies.

## Introduction

Malaria remains a major cause of mortality and morbidity in Africa with close to 600,000 deaths reported in 2023 (1). Implementation of vector control tools reliant on insecticides, including insecticide treated bed nets (ITNs) and indoor residual spraying (IRS) have largely contributed in reducing the disease incidence (2), but their power and sustainability is threatened by the selection of high levels of insecticide resistance (3, 4).

The last decade a wealth of studies has identified and characterized genes in the main malaria vector *Anopheles gambiae* with a role in insecticide resistance using transcriptomic, genomic and molecular approaches (5–7). Two main categories of resistance mechanisms have been described: the toxicokinetic, that involve mutations at the insecticide target sites that reduce binding efficiency, and the toxicodynamic, that reduce the amount of insecticide reaching its target, either through increased metabolism and sequestration or decreased penetration and increased excretion (8).

Among the most well-known toxicokinetic cases are mutations in the pyrethroid target site, the voltage gated sodium channel (VGSC), with the classical L995F (widely known as L1014F) mutation being widespread in field populations, often reaching fixation (9). This mutation has been previously functionally validated through CRISPR editing showing to confer up to 14-fold resistance to pyrethroids (10).

In the case of toxicodynamic mechanisms, cytochrome P450s have been characterized as a key enzyme family involved in detoxification of xenobiotics through hydroxylation (Phase I metabolism) (11, 12). In *Anopheles* mosquitoes, over-expression of members of the CYP6 and CYP9 families, including *Cyp6m2*, *Cyp6p3* and *Cyp9k1* are commonly over-expressed in resistant populations and have been shown to metabolise pyrethroids (12–14). Glutathione *S*-transferases have also been associated with insecticide resistance. They can act either directly by adding a glutathione molecule to their substrate insecticide, rendering it more water soluble (Phase II metabolism) or indirectly through sequestration and/or reduction of oxidative stress (15). *Gste2* has been shown to confer resistance to DDT and fenitrothion in *An. gambiae*, while it has also been associated with resistance to permethrin (16, 17). Carboxyl-esterases are another type of detoxification enzyme that act primarily through sequestration of their substrate. Recently Coeae6g was functionally validated showing to confer resistance to the organophosphate pirimiphos-methyl and permethrin (18). Increased excretion of insecticides has also been described as a resistance mechanism (Phase III), but is less well characterized. Evidence was provided previously for the association of the ABCH2 transporter in deltamethrin toxicity, by acting as a pyrethroid exporter located in the legs (19).

Although there are cases in mosquitoes where a single mechanism has been described to drive resistance, as is the case of mutations on the chitin synthase in *Culex pipiens,* conferring high levels of resistance to diflubenzuron (20) and the over-expression of *Cyp6P9a/b* in *An. funestus* conferring pyrethroid resistance (21), in most cases resistance has a complex genetic basis where multiple genes are associated with the phenotype (12, 22–24). *An. gambiae* field populations frequently harbor target site resistance mutations alongside the overexpression of various detoxification enzymes, from the same and/or different enzyme families (25). In those more complex cases it is important to understand how different mechanisms combine to shape resistance levels. Are individual mechanisms sufficient to confer resistance and to what extent? Moreover, does the co-existence of different mechanisms result in additive or even synergistic effects that would explain their persistence in the same genetic background? Gaining this knowledge is essential for understanding the dynamics of insecticide resistance evolution and designing efficient resistance surveillance and management strategies.

In this study, we used a large panel of transgenic strains and crosses between them to validate the role of multiple detoxification enzymes, including the P450s CYP6P3, CYP6M2, CYP9K1; the carboxyl esterase COEAE6G; the glutathione S-transferase GSTE2 and the ABC transporter ABCH2, when alone or in combination with each other or the *kdr* target site mutation, in conferring pyrethroid resistance and its magnitude. We also used the P450 over-expressing strains to study the phenotypic responses to pro-insecticide exposure. Pro-insecticides require activation to their more toxic form by endogenous P450s, and thus are considered a good option for managing resistant populations with elevated P450 expression. However, as P450s can also detoxify pro-insecticides, it is critical to know if their activation or detoxification will prevail *in vivo,* especially in cases were *in vitro* metabolism assays provide inconclusive results (26). Here we test cross resistance to the mitochondrial uncoupler chlorfenapyr, a new generation malaria preventative product that is used in dual insecticide bednets, the sodium channel blocker indoxacarb and the organophosphate pirimiphos-methyl, that is increasingly being used in IRS applications (27, 28).

## Results

### Functional validation of resistance levels conferred by the over-expression of Abch2 alone and in combination with Cyp6p3

We used the GAL4-UAS system to over-express *Abch2* in a susceptible lab strain and validate its role in resistance functionally. Initially a UAS-Abch2 responder line marked with YFP (under the 3xP3 promoter) was generated through site-directed recombination-mediated cassette exchange (RMCE) into the CFP marked (under the 3×P3 promoter) A11 docking line (2×attP, generated in Adolfi et al., 2018 (14)) (Supplementary Figure 1A). Seven F1 progeny were identified with cassette exchange (marked with YFP only) and used for the establishment of the line (details provided in Supplementary table 2). The UAS-Abch2 line was crossed to a GAL4 driver line that expresses the trans-activator under the ubiquitin promoter (Ubi-GAL4), driving expression in multiple tissues (line generated and described in Adolfi et al.,2018 (14, 29)). Progeny of this cross transcribed *Abch2* (hereafter marked Abch2^+^) at approximately 66-fold (Supplementary Figure 2A) higher levels compared to the A11 parental line.

To assess how the Abch2 over-expression impacts the susceptibility of mosquitoes to insecticides we used female Abch2^+^ and exposed them to WHO tube toxicity bioassays. WHO tube assays are used to assess the presence of resistance by exposing mosquitoes to discriminating doses (twice the lethal concentration that kills 99% of a susceptible population). Mortality (24 h post exposure) at the discriminating dose of less than 90% indicates resistance (30). For all insecticides tested, including the pyrethroids: deltamethrin, permethrin and *α*-cypermethrin; the organophosphates: malathion and pirimiphos-methyl, the carbamate: bendiocarb and the organochlorine: DDT, no resistance was observed for Abch2^+^ mosquitoes (100% mortality, 24h post exposure) (Supplementary Figure 3A). However, a significantly lower knock down rate (defined as the inability to stand up within an hour of exposure) compared to the control A11 group was observed for the three pyrethroids (Supplementary Figure 3B).

To investigate further whether Abch2 provides reduced sensitivity to pyrethroids, we used WHO papers impregnated with 0.016% deltamethrin, instead of the 0.05% discriminating dose, as previously used by Kefi et al., 2023 (19). We also varied the exposure time, to estimate the Lethal time 50 (LT_50_) and quantify sensitivity levels. Significantly lower mortality was observed for Abch2^+^ mosquitoes compared to control mosquitoes at all time points tested (Figure 1A). The LT_50_ of the Abch2^+^ was estimated at 25 min. Given that at 3 min exposure, mortality of the A11 was around 50%, the *Abch2* over-expressing mosquitoes show a minimum resistance ratio of 8-fold at the 0.016% deltamethrin dose (Figure 1A and Supplementary Table 3).

**Figure 1.**
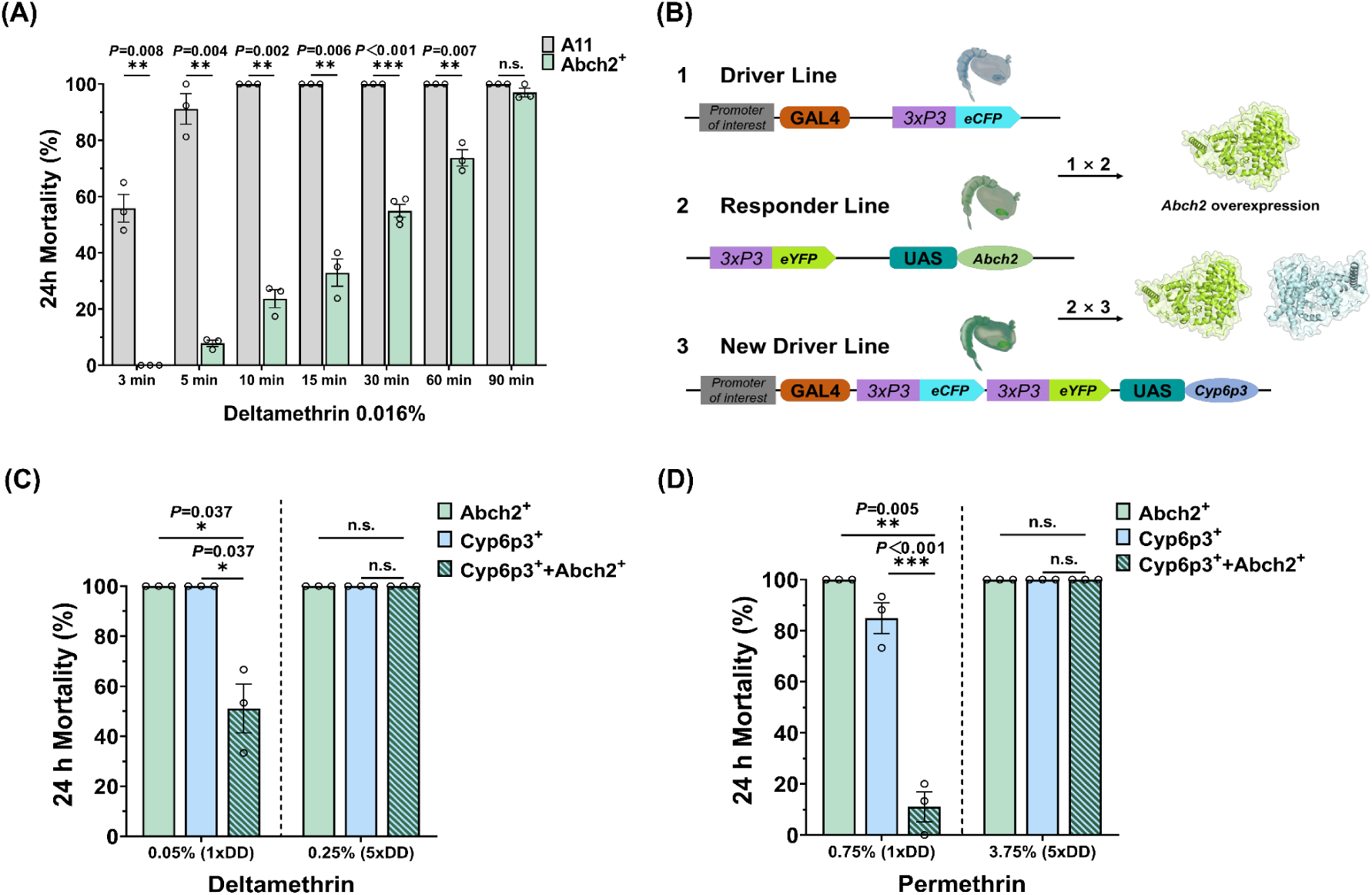
Evaluating the effect of multi-tissue over-expression of *Abch2*, when alone and in combination with *Cyp6p3* on pyrethroid sensitivity. (A) Percentage mortality of Abch2⁺ and control (A11) mosquitoes 24 h after exposure to 0.016% deltamethrin impregnated papers in a WHO tube assay across different exposure times. (B) Schematic diagram showing the UAS-Abch2 responder and the Ubi-GAL4: UAS-Cyp6p3 lines and how the GAL4/UAS system drives overexpression of the target genes either individually or in combination. (C) Percentage mortality of mosquitoes over-expressing *Abch2* (Abch2^+^), *Cyp6p3* (Cyp6p3^+^) or their combination (Abch2⁺+Cyp6p3^+^), 24 h after exposure to 1× (0.05%) and 5× (0.25%) WHO diagnostic dose of deltamethrin (D) Percentage mortality of Abch2^+^, Cyp6p3^+^ and Abch2⁺+Cyp6p3^+^ mosquitoes, 24 h after exposure to 1× (0.75%) and 5× (3.75%) WHO diagnostic dose of permethrin. For all bioassays, at least three biological replicates were performed, each comprising 20–25 3∼5-day-old females. Individual data points represent different biological replicates, and bars indicate the standard error (SE). Statistical significance was assessed using Welch’s *t*-test (n.s., not significant, \**P*< 0.05, \*\**P* < 0.01; \*\*\**P* < 0.001).

To assess whether there is a combined effect of ABCH2 and P450 over-expression, we co-over-expressed the transporter with *Cyp6p3*. To do so we initially generated a line combining at the same locus the Ubi-GAL4 and the UAS-Cyp6p3 elements, thus natively over-expressing *Cyp6p3,* without the need of crossing. To generate this line, we used the Ubi-GAL4 line as a docking line and integrated the UAS-Cyp6p3 cassette (individuals marked YFP and CFP), as depicted in Supplementary figure 4A and described by Adolfi et al., 2019 (14). The generated Ubi-GAL4: UAS-Cyp6p3 line (in short hereafter GAL4-Cyp6p3^+^) natively over-expressed *Cyp6p3* at 401-fold (transgene at one copy), which is lower than the 573-fold over-expression of *Cyp6p3* seen in progeny of the Ubi-GAL4 x UAS-Cyp6p3 (strain previously generated by Adolfi et al., 2019) cross, but shows similar resistance levels (Supplementary figure 4B-D). The GAL4-Cyp6p3^+^ strain was crossed to the UAS-Abch2 and progeny (Abch2^+^+ Cyp6p3^+^) were tested for the over-expression of both genes (122-fold for *Cyp6p3* and 21-fold for *Abch2*) (Supplementary figure 2D).

Abch2^+^+Cyp6p3^+^ mosquitoes showed significantly lower mortality to the 1× diagnostic dose of deltamethrin and permethrin compared to mosquitoes over-expressing only one of the two genes (Figure 1B, C). To assess the strength of resistance, exposure to 5× and 10× diagnostic doses is recommended by WHO. Abch2^+^+Cyp6p3^+^ mosquitoes showed 100% mortality at the 5× diagnostic dose (Figure 1 B, C).

### Functional validation of resistance levels conferred by the over-expression of Cyp9k1 alone and in combination with Cyp6p3

A UAS-Cyp9k1 responder line, marked with YFP (under the 3×P3 promoter) was generated as described in the previous section for the UAS-Abch2 strain. Details of the establishment of the strain are provided in Supplementary Figure 1B and Supplementary Table 2. Progeny of the cross between UAS-Cyp9k1 and the driver line Ubi-GAL4 (in short Cyp9k1^+^) over-expressed *Cyp9k1* 19-fold (Supplementary Figure 2B).

Mosquitoes over-expressing *Cyp9k1* were exposed to WHO discriminating doses of several insecticides. Resistance was observed for the pyrethroids deltamethrin (67% mortality), permethrin (19% mortality) and *α*-cypermethrin (18% mortality) (Figure 2). A smaller, but significant reduction in mortality to DDT (85%) was also observed. No resistance was observed for bendiocarb, malathion and pirimiphos-methyl (Figure 2). We produced time response curves to quantify LT_50_ resistance levels of Cyp9k1^+^ mosquitoes. The resistance ratio was estimated at ≥ 12-fold for deltamethrin, 11-fold for permethrin, ≥ 29-fold for *α*-cypermethrin and 2-fold for DDT (Table 1).

**Figure 2.**
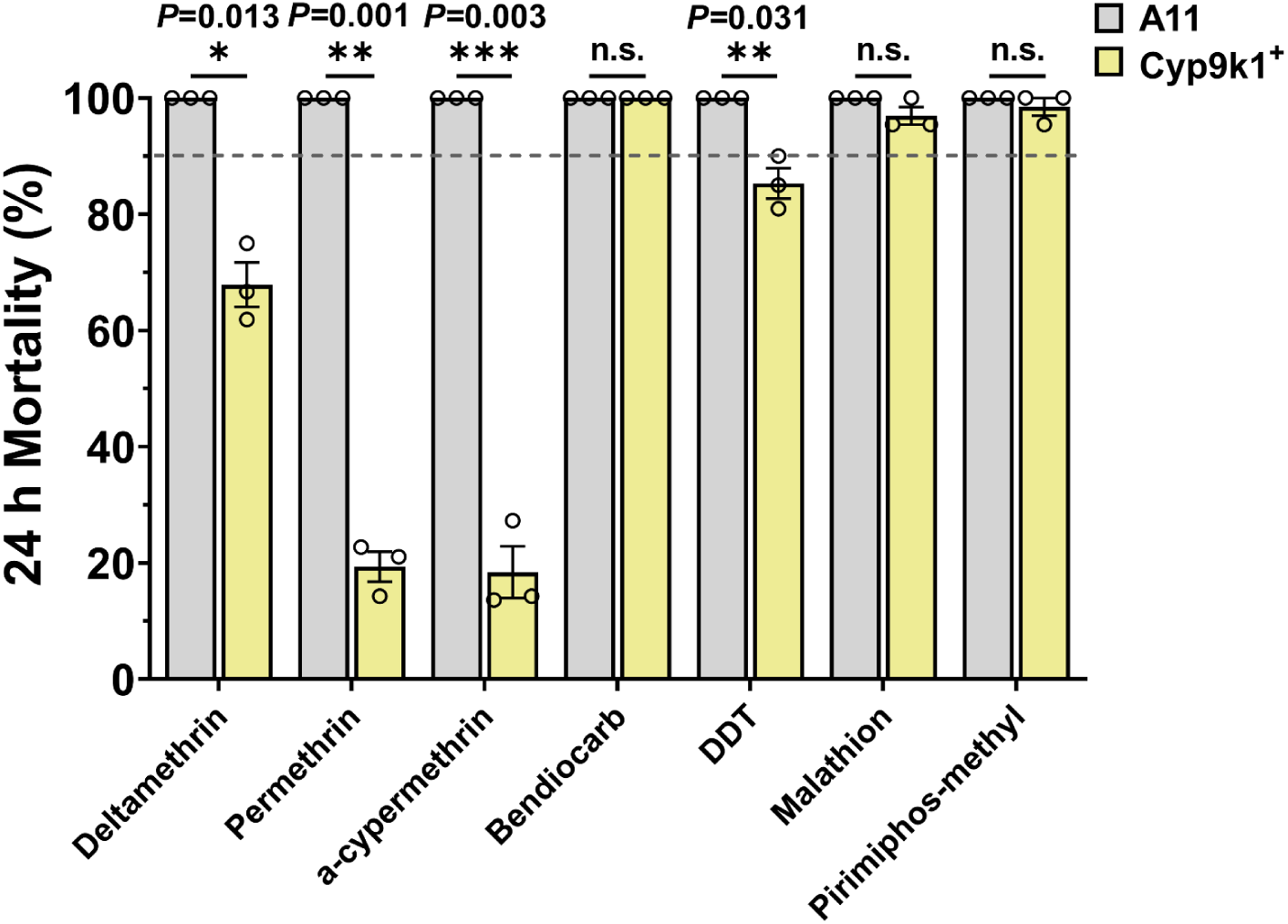
WHO toxicity bioassays for testing the effect of multi-tissue *Cyp9k1* over-expression on insecticide resistance. Percentage mortality of *Cyp9k1* over-expressing (Cyp9k1^+^) and control (A11) mosquitoes 24 h after 1h exposure to the 1× WHO diagnostic dose of different insecticides (0.05% deltamethrin, 0.75% permethrin, 0.05% *α*-cypermethrin, 0.1% bendiocarb, 4% DDT, 5% malathion, and 0.25% pirimiphos-methyl). For all bioassays, at least three biological replicates were performed, each comprising 20–25 randomly selected 3∼5-day-old females. Individual data points represent different biological replicates, and bars indicate the standard error (SE). The dotted line indicates the WHO 90% mortality threshold used to define resistance. Statistical significance was assessed using Welch’s *t*-test (n.s., not significant, \**P*< 0.05, \*\**P* < 0.01; \*\*\**P* < 0.001).

**Table 1.** Time-response toxicity assays for estimating resistance strength of *Cyp9k1* over-expressing mosquitoes. Cyp9k1^+^ and A11 control mosquitoes were exposed to the 1× diagnostic dose of insecticides in WHO tube assays varying the exposure time. The LT_50_ (time required to obtain 50% mortality) is provided for each strain with the 95% fiducial limits (95% FL) and used for calculating the resistance ratio (RR) (LT_50_ of Cyp9k1^+^ / LT_50_ of A11). Each bioassay included at least five exposure time points, for each of which at three replicates, of 20–25, 3∼5-day-old female mosquitoes were tested. **\*** A11’s LT_50_ value for permethrin was obtained from a previous publication using the same strain(18).

| Line | Insecticides | LT<sub>50</sub> | 95%FL | Slope±SE | RR |
| --- | --- | --- | --- | --- | --- |
| A11 | Deltamethrin | <3 |  |  |  |
|  | Permethrin | 7.3* |  |  |  |
|  | α-Cypermethrin | <3 |  |  |  |
|  | DDT | 19.74 | 17.56 to 22.25 | 3.63±0.36 |  |
| Cyp9k1<sup>+</sup> | Deltamethrin | 38.06 | 32.71 to 43.56 | 3.10±0.31 | >12.67 |
|  | Permethrin | 81.46 | 69.89 to 93.98 | 4.34±0.41 | 11.16 |
|  | α-Cypermethrin | 89.82 | 79.12 to 101.83 | 4.49±0.56 | >29.94 |
|  | DDT | 47.06 | 43.09 to 51.43 | 8.94±0.80 | 2.39 |

Next, we assessed whether there is a combined effect between *Cyp9k1 and Cyp6p3* over-expression. We crossed the GAL4-Cyp6p3^+^ strain with the UAS-Cyp9k1 responder line and measured the over-expression levels of both genes in their progeny (Cyp9k1^+^+Cyp6p3^+^): 26-fold for *Cyp9k1* and 125-fold for *Cyp6p3* (Supplementary figure 2E). Resistance, and its strength to deltamethrin and permethrin were evaluated through WHO tube bioassays using 1×, 5× and 10× diagnostic doses. Mosquitoes over-expressing both P450s showed resistance not only to 1× deltamethrin and 1× permethrin, but also to the 5× deltamethrin and 5× permethrin diagnostic dose with significantly lower mortality compared to mosquitoes over-expressing only one of the two genes (Figure 3A and 3B). 100% mortality was observed in the 10× diagnostic doses for all strains (Figure 3A and 3B).

**Figure 3.**
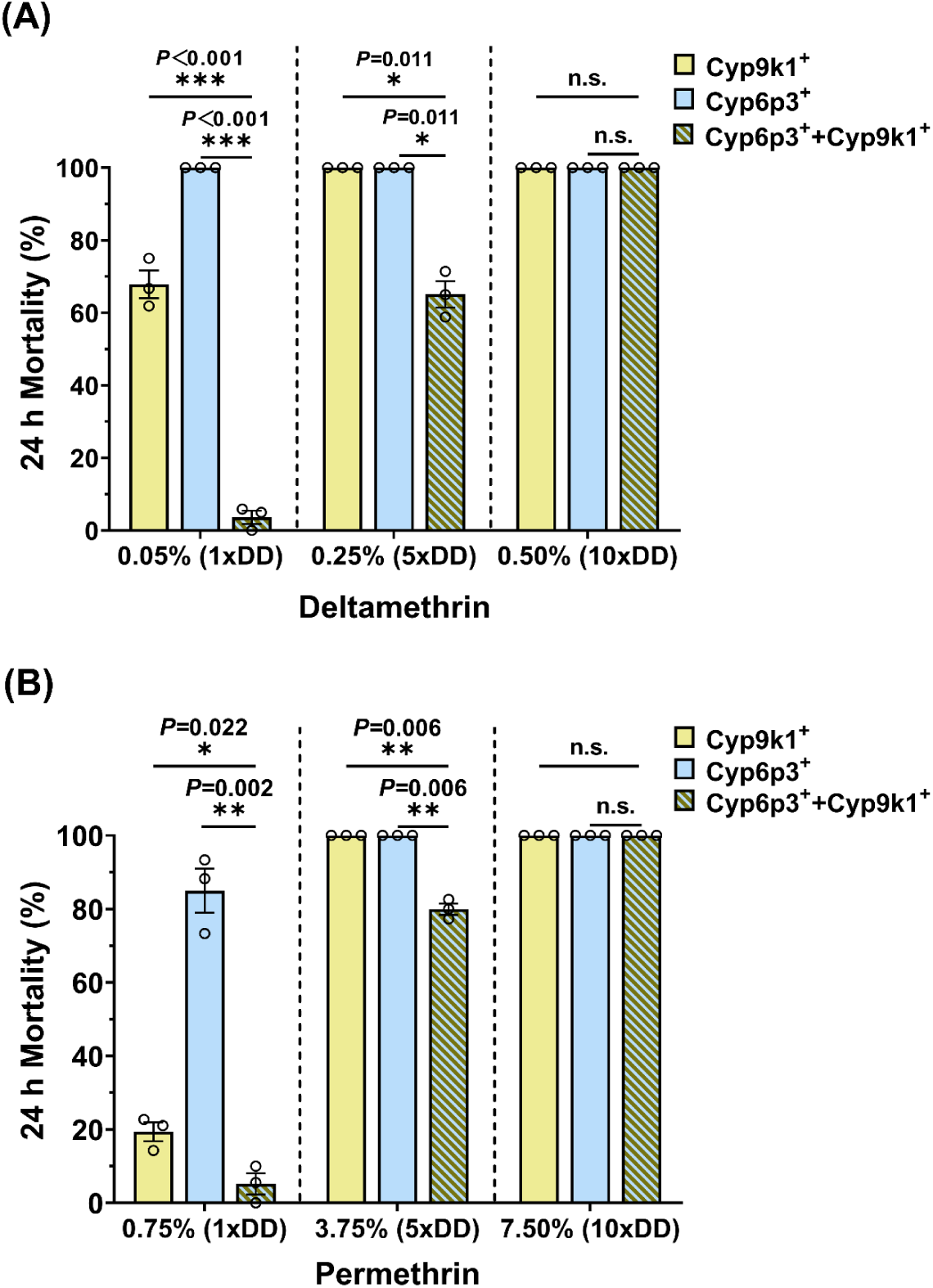
Evaluating the effect of *Cyp9k1* and *Cyp6p3* co-overexpression on pyrethroid resistance levels. (A) Percentage mortality of mosquitoes over-expressing *Cyp9k1* (Cyp9k1^+^), *Cyp6p3* (Cyp6p3^+^) or their combination (Cyp9k1^+^+Cyp6p3^+^) 24 h after 1h exposure to the 1× (0.05%), 5× (0.25%) and 10× (0.50%) WHO diagnostic dose of deltamethrin. (B) Percentage mortality of Cyp9k1^+^, Cyp6p3^+^ and Cyp9k1^+^+Cyp6p3^+^ mosquitoes 24 h after exposure to the 1× (0.75%), 5× (3.75%) and 10× (7.50%) WHO diagnostic dose of permethrin. For all bioassays, at least three replicates were performed, each comprising 20–25 3∼5-day-old females. Individual data points represent different replicates, and bars indicate the standard error (SE). Statistical significance was assessed using Welch’s *t*-test (n.s., not significant, \**P*< 0.05, \*\**P* < 0.01; \*\*\**P* < 0.001).

To test if the observed effect is due to higher expression of pyrethroid metabolizing P450s or higher expression of the specific two P450s, we crossed the GAL4-Cyp6p3^+^ strain to the responder line UAS-Cyp6p3 and assessed pyrethroid resistance of their progeny (Cyp6p3^+^+Cyp6p3^+^). Progeny showed 898- fold over-expression of *Cyp6p3* (Supplementary Figure 4B), and low or no resistance to the 1×, 5× and 10× diagnostic dose of deltamethrin and permethrin (Supplementary Figure 4C, D).

### Evaluating the combined effect of two different type detoxification enzymes

We next tested the combined effect of two enzymes belonging to different detoxification enzyme families. We combined the over-expression of *Cyp6p3* with that of *Gste2* and of the carboxylesterase *Coeae6g*. To do that we crossed the GAL4-Cyp6p3^+^ strain with previously generated UAS-Gste2 and UAS-Coeae6g responder lines(14, 18). From those crosses we obtained progeny over-expressing *Cyp6p3* and *Gste2* (Cyp6p3^+^+Gste2^+^) and progeny over-expressing *Cyp6p3* and *Coeae6g* (Cyp6p3^+^+Coeae6g^+^) (Supplementary Figure 2F, G) respectively. We also crossed Ubi-GAL4 separately with the UAS-Cyp6p3, UAS-Gste2 and UAS-Coeae6g lines (Coeae6g^+^) to obtain progeny that over-express only one gene. Cyp6p3^+^+Gste2^+^ mosquitoes were susceptible to deltamethrin, having the same response as the Cyp6p3^+^ and Gste2^+^ mosquitoes (Figure 4A), while for permethrin they showed a small but significant decrease in mortality compared to the Cyp6p3^+^ and Gste2^+^ mosquitoes in the 1× diagnostic dose. The Cyp6p3^+^+Coeae6g^+^ progeny were tested for permethrin and pirimiphos-methyl, two insecticides that have been shown to interact *in vitro* with both CYP6P3 and COEAE6G(18, 26). The Cyp6p3^+^+Coeae6g^+^ mosquitoes showed significantly lower mortality compared to the strains expressing one of the two enzymes when exposed to 1x and 5x diagnostic doses of permethrin (Figure 4C), while for pirimiphos-methyl they had approximately the same response as the Coeae6g^+^ mosquitoes (Figure 4D)(18).

**Figure 4.**
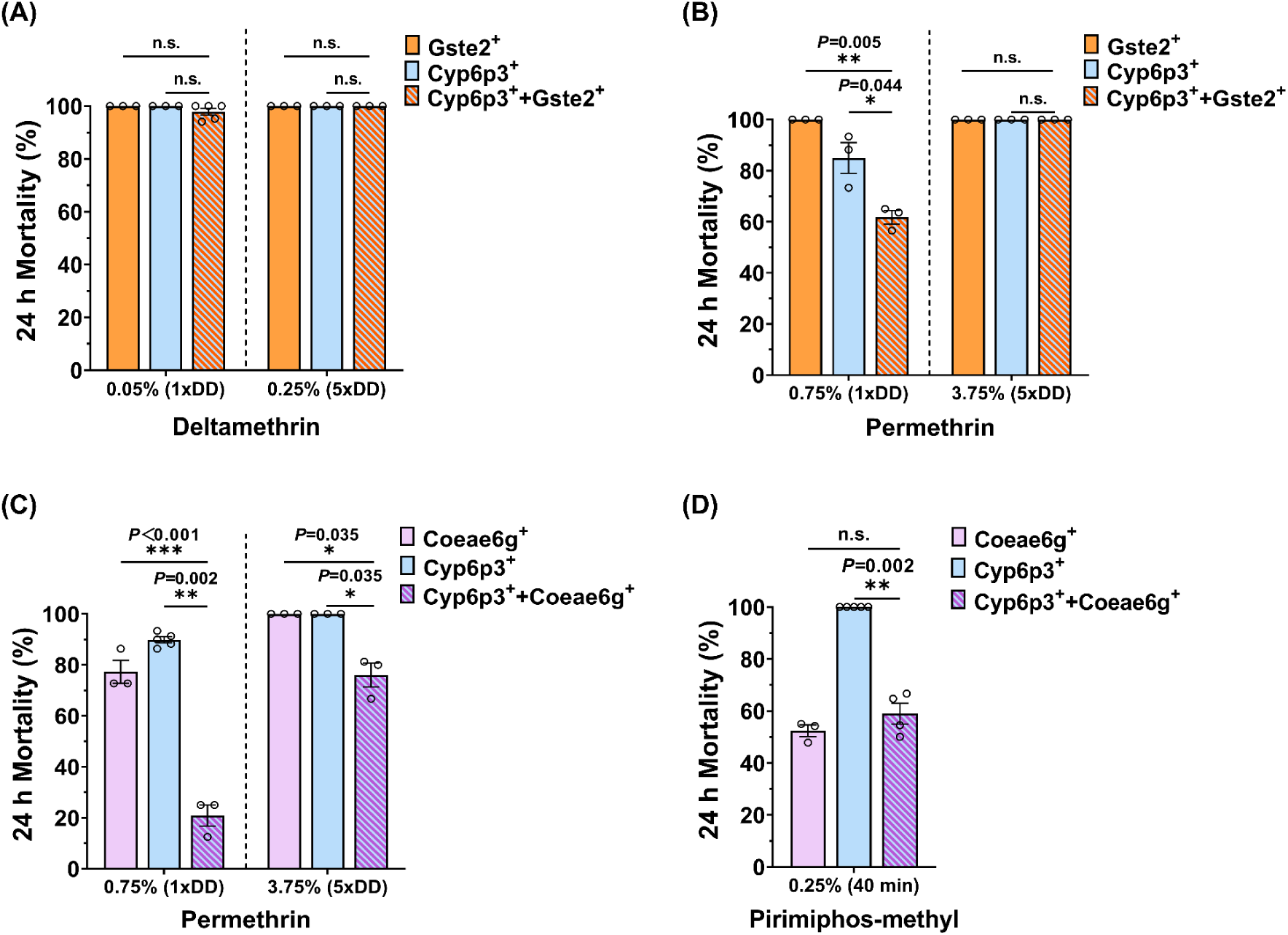
Evaluating the effect of *Cyp6p3* co-overexpression with *Gste2* or *Coeae6g* on insecticide resistance levels. (A) Percentage mortality of mosquitoes over-expressing *Gste2* (Gste2^+^), *Cyp6p3* (Cyp6p3^+^) or their combination (Gste2^+^+Cyp6p3^+^) 24 h after 1h exposure to the 1× (0.05%) and 5× (0.25%) WHO diagnostic dose of deltamethrin and (B) 1h exposure to the 1× (0.75%) and 5× (3.75%) WHO diagnostic dose of permethrin. (C) Percentage mortality of mosquitoes over-expressing *Coeae6g* (Coeae6g^+^), *Cyp6p3* (Cyp6p3^+^) or their combination (Coeae6g^+^+Cyp6p3^+^) 24 h after 1h exposure to the 1× (0.75%) and 5× (3.75%) WHO diagnostic dose of permethrin and (D) 40min exposure the 1× (0.25%) diagnostic dose of pirimiphos-methyl, which is the previously estimated LT_50_ for Coeae6g^+^. For all bioassays, at least three replicates were performed, each comprising 20–25 3∼5-day-old females. Individual data points represent different replicates, and bars indicate the standard error (SE). Statistical significance was assessed using Welch’s *t*-test (n.s., not significant, \**P*< 0.05, \*\**P* < 0.01; \*\*\**P* < 0.001).

### Evaluating the combined effect of target site and metabolic resistance

We also tested the combined effect of metabolic and target site resistance mechanisms by combining the VGSC mutation L995F (hereafter called *kdr*) with the over-expression of *Cyp6p3* and *Cyp6m2*. We introduced the *kdr* mutation in the driver (Ubi-GAL4) and responder lines (UAS-Cyp6p3 and UAS-Cyp6m2) through crosses with a previously generated CRISPR edited line, that carries the 995F mutation in an insecticide susceptible genetic background(10). More details on the establishment of the Ubi-GAL4+*kdr*, UAS-Cyp6p3+*kdr* and UAS-Cyp6m2+*kdr* lines are provided in Supplementary Figure 5A-C. By crossing the Ubi-GAL4+*kdr* line with the two responder lines UAS-Cyp6p3+*kdr* and UAS-Cyp6m2+*kdr* we obtained progeny that carry the *kdr* mutation in homozygosity and over-expressed the *Cyp6p3* (CYP6p3^+^+*kdr*) or *Cyp6m2* (CYP6m2^+^+*kdr),* respectively (Supplementary Figure 5D, E).

The combination of *kdr* with over-expression of *Cyp6p3* conferred high levels of resistance to deltamethrin in all: 1×, 5× and 10× diagnostic doses and significantly lower mortality compared to the strains carrying only one of the two mechanisms (Figure 5A). The combination of *kdr* mutation with the over-expression of *Cyp6m2,* conferred resistance and significantly lower mortality compared to the strains carrying one of the two mechanisms, only in the case of the 1× diagnostic dose (Figure 5B). We also generated time response curves to deltamethrin and estimated the LT_50_ values for all strains (Table 2). As shown in Table 2 the LT_50_ values for the Cyp6p3^+^+*kdr* and Cyp6m2^+^+*kdr* strains are greater than the sum of the LT_50_ values of the individual mechanisms. Thus, the combination of the two mechanisms acts synergistically, conferring high levels of resistance.

**Figure 5.**
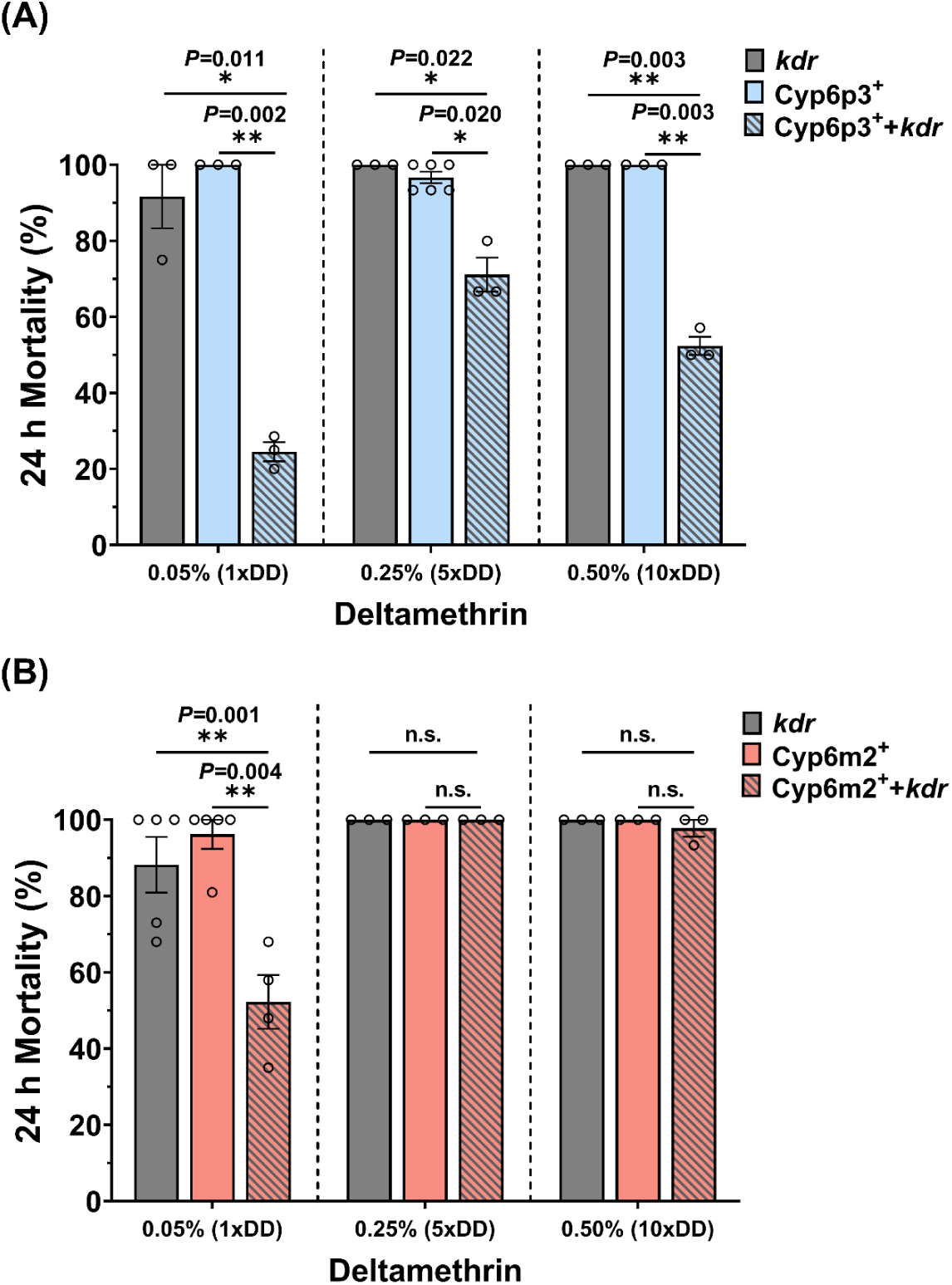
Evaluating the combined effect of *Cyp6p3* or *Cyp6m2* over-expression with the VGSC mutation 995F (*kdr*) on pyrethroid resistance levels. (A) Percentage mortality of mosquitoes: over-expressing *Cyp6p3* (Cyp6p3^+^), carrying the L995F mutation (*kdr*) or having the combination of the two mechanisms (Cyp6p3^+^+*kdr*) 24h after 1h exposure to 1× (0.05%), 5× (0.25%) and 10× (0.50%) WHO diagnostic dose of deltamethrin. (B) Percentage mortality of mosquitoes: over-expressing *Cyp6m2* (Cyp6m2^+^), carrying the L995F mutation (*kdr*) or having the combination of the two mechanisms (Cyp6m2^+^+*kdr*) 24 h after 1h exposure to 1× (0.05%), 5× (0.25%) and 10× (0.50%) WHO diagnostic dose of deltamethrin. For all bioassays, at least three replicates were performed, each comprising 20–25, 3∼5-day-old females. Individual data points represent different biological replicates, and bars indicate the standard error (SE). Statistical significance was assessed using Welch’s *t*-test (n.s., not significant, \**P*< 0.05, \*\**P* < 0.01; \*\*\**P* < 0.001).

**Table 2.** Time-response toxicity assays for estimating resistance strength in mosquitoes carrying combinations of target site and metabolic resistance. Mosquitoes carrying the VGSC L995F mutation (*kdr*), over-expressing one of the two P450s (*Cyp6p3*, *Cyp6m2*) or having a combination of the two mechanisms (Cyp6p3^+^+*kdr*, Cyp6m2^+^+*kdr*) were exposed to the 1× WHO diagnostic dose of deltamethrin at varying exposure times. The LT_50_ (time required to obtain 50% mortality) is provided for each strain with the 95% fiducial limits (95% FL). The resistance ratio (RR) (LT_50_ of resistant strain/ LT_50_ of control), is given in comparison with the A10 control strain. For each LT_50_ a minimum of five exposure time points were tested with at least three replicates per time point. Each replicate, includes 20–25 3∼5-day-old female mosquitoes.

| Line | Insecticides | LT<sub>50</sub> | 95%FL | Slope±SE | RR |
| --- | --- | --- | --- | --- | --- |
| A10 | Deltamethrin | 2.99 | 1.93-4.06 | 3.11±0.30 | - |
| *kdr* | Deltamethrin | 18.92 | 13.44-25.20 | 2.57±0.24 | 6.33 |
| *Cyp6p3*<sup>+</sup> | Deltamethrin | 11.35 | 8.50-13.98 | 2.56±0.23 | 3.80 |
| *Cyp6m2*<sup>+</sup> | Deltamethrin | 7.26 | 4.56-10.66 | 1.613±0.15 | 2.48 |
| *Cyp6p3*<sup>+</sup>*kdr* | Deltamethrin | 174.63 | 129.95-305.53 | 2.23±0.30 | 58.40 |
| *Cyp6m2*<sup>+</sup>*kdr* | Deltamethrin | 79.65 | 54.33-112.73 | 1.97±0.19 | 26.63 |

### Using transgenic mosquitoes over-expressing P450s to assess negative cross resistance to pro-insecticides

Initially we gathered published data on metabolic activity of CYP6P3 and CYP9K1, in the form of substrate depletion and formation of metabolites, towards the pro-insecticides pirimiphos-methyl, indoxacarb and chlorfenapyr (Table 3)(26, 31). As there were no data on the metabolism of indoxacarb by CYP6P3 and of pirimiphos-methyl by CYP9K1 we supplemented those by recombinantly expressing the two proteins and using them in metabolism assays. Both enzymes have been shown to metabolize chlorfenapyr to the active form tralopyril, while no other metabolite was detected by mass spectrometry with selected ion monitoring(31). Both enzymes also metabolize pirimiphos-methyl (here we show a 35% depletion of pirimiphos-methyl by CYP9K1) and generate two metabolites: the active pirimiphos-methyl oxon and the oxidatively cleaved 2-diethylamino-6-hydroxyl-4-methylpyrimidine, with similar abundance (26) (Supplementary Figure 8A-D). Indoxacarb is also metabolized by both enzymes (here we show a 19,5% depletion by CYP6P3 and 60% by CYP9K1). In the case of CYP6P3 a hydroxylated metabolite was identified. However, the exact place of the hydroxylation could not be determined and thus the metabolite remains uncharacterized (Supplementary Figure 6A-D). In the case of CYP9K1 both the active DCJW and a hydroxylated metabolite (again uncharacterized) were identified (Supplementary Figure 7A-E).

**Table 3.** Metabolic activity of recombinantly expressed CYP6P3 and CYP9K1 for chlorfenapyr, Indoxacarb and pirimiphos-methyl. Percentage depletion of each insecticide measured by HPLC and formation of metabolites measured with mass spectrometry.

| Protein | Insecticides | Depletion | Identified metabolites | Note |
| --- | --- | --- | --- | --- |
| **CYP6P3** | Indoxacarb | 19.5% | ● Hydroxylated (uncharacterized) | This study |
|  | Chlorfenapyr | >80% | ● Tralopyril (active) | Yunta et al., 2023(31) |
|  | Pirimiphos-methyl | 100% | ● Pirimiphos methyl-oxon (active)<br>● Oxidatively cleaved: 2-diethylamino-6-hydroxyl-4-methylpyrimidine | Yunta et al., 2019(26) |
| **CYP9K1** | Indoxacarb | 60% | ● DCJW (active)<br>● Hydroxylated (uncharacterized) | This study |
|  | Chlorfenapyr | >90% | ● Tralopyril (active) | Yunta et al., 2023(31) |
|  | Pirimiphos-methyl | 35.9% | ● Pirimiphos methyl-oxon (active)<br>● Oxidatively cleaved (2-diethylamino-6-hydroxyl-4-methylpyrimidine) | This study |

To test *in vivo* the balance between activation or detoxification of the pro-insecticides, we used the Cyp9k1^+^ and Cyp6p3^+^ strains and exposed them to the pro-insecticides through topical bioassays, which allow more precise insecticide doses to be administered. Both strains showed ∼10-fold resistance to indoxacarb (compared to the A11 control strain). Cyp6p3^+^ was 1.9-fold more susceptible to chlorfenapyr and 1.7-fold to pirimiphos-methyl. Cyp9k1^+^ was 2.5-fold more susceptible to chlorfenapyr and 2.4-fold to pirimiphos-methyl (Table 4). We also tested resistance levels to the pro-insecticides when combining the over-expression of *Cyp6p3* and *Cyp9k1*. Over-expression of both genes (expression levels shown in Supplementary figure 2E) resulted in small but significant changes in the sensitivity to indoxacarb and chlorfenapyr compared to Cyp9k1^+^ mosquitoes but not compared to Cyp6p3^+^ (Figure 6). No difference was observed in PM sensitivity by co-expression.

**Figure 6.**
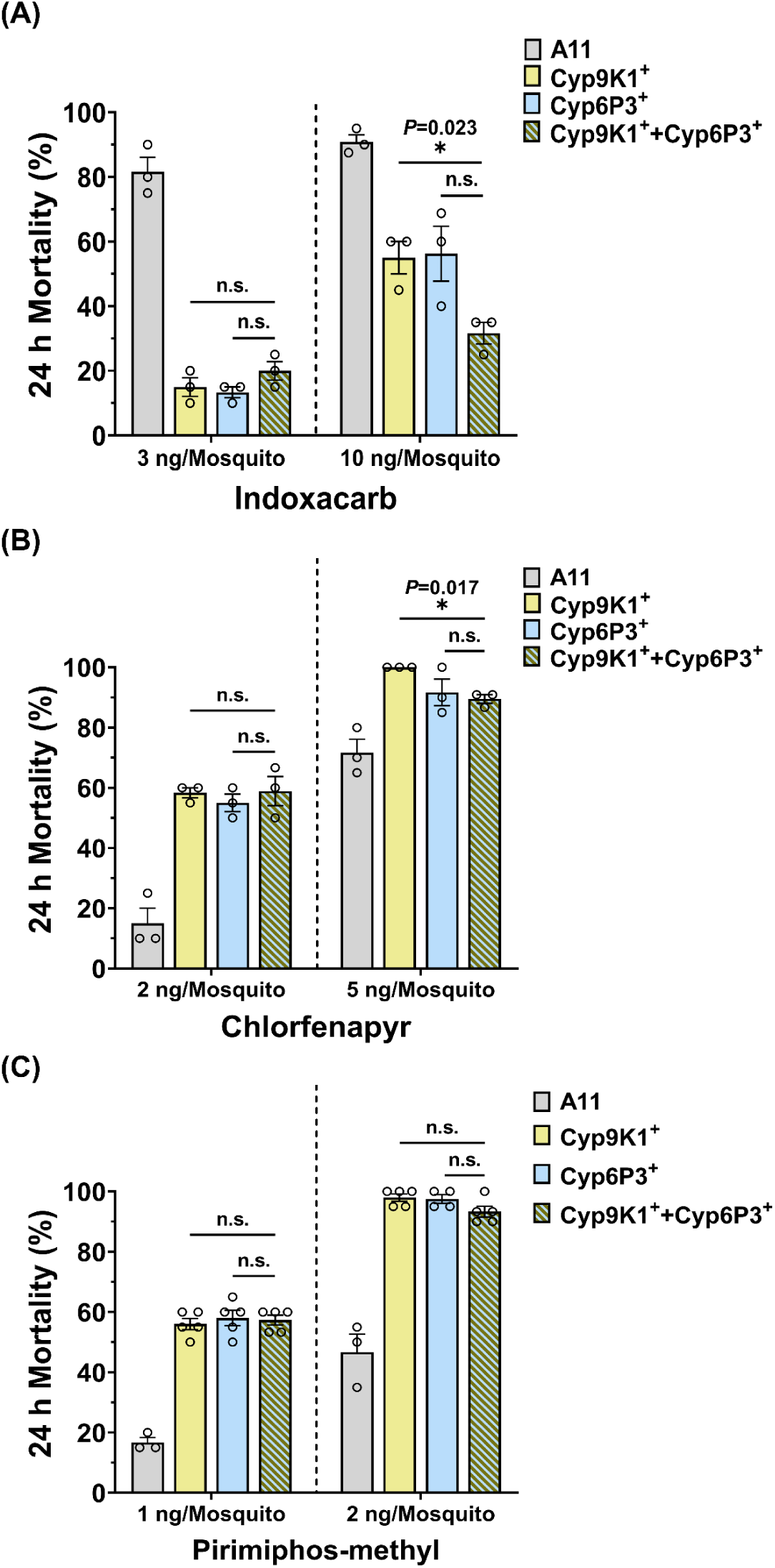
Evaluating the effect of combined *Cyp6p3* and *Cyp9k1* overexpression on the sensitivity to pro-insecticides. (A) Percentage mortality of mosquitoes over-expressing *Cyp9k1* (Cyp9k1^+^), *Cyp6p3* (Cyp6p3^+^) or their combination (Cyp9k1^+^+Cyp6p3^+^) 24 h after a topical application of 3 and 10 ng of indoxacarb per mosquito, that are equivalent to the LD_20_ and LD_50_ of the Cyp6p3^+^strain (B) 2 and 5 ng chlorfenapyr per mosquito, that are equivalent to the LD_50_ and LD_90_ of the Cyp6p3^+^ strain, (C) 1 and 2 ng pirimiphos-methyl per mosquito, equivalent to the LD_50_ and LD_90_ of the Cyp6p3^+^ strain. For all bioassays, at least three replicates were performed, each comprising 20 3∼5-day-old females. Individual data points represent different replicates, and bars indicate the standard error (SE). Statistical significance was assessed using Welch’s *t*-test (n.s., not significant, \**P*< 0.05, \*\**P* < 0.01; \*\*\**P* < 0.001).

**Table 4.** Evaluating sensitivity to pro-insecticides in Cyp6p3^+^ and Cyp9k1^+^ mosquitoes through dose response toxicity assays. Toxicity bioassays through topical application of insecticides were performed in mosquitoes over-expressing *Cyp6p3* and *Cyp9k1*, as well as control mosquitoes (A11). The LD_50_ (concentration causing 50% mortality) with 95% fiducial limits is provided for each strain. Resistance ratio (RR) is provided in comparison to A11 (LD_50_ of Cyp6p3^+^/Cyp9k1^+^ divided by LD_50_ of A11). For the estimation of LT_50_ at least five concentrations were used, each including at least three biological replicates. For each replicate, twenty 3∼5-day-old female mosquitoes were tested.

| Line | Insecticides | LD<sub>50</sub><br>(ng/mosquitoes) | 95%FL | Slope±SE | RR |
| --- | --- | --- | --- | --- | --- |
| **A11** | Indoxacarb | 1.12 | 0.69 to 1.58 | 1.42±0.14 |  |
|  | Chlorfenapyr | 3.82 | 3.21 to 4.66 | 3.14±0.38 |  |
|  | Pirimiphos-methyl | 1.71 | 1.34 to 2.22 | 2.61±0.23 |  |
| **Cyp6p3<sup>+</sup>** | Indoxacarb | 12.49 | 10.27 to 15.14 | 1.76±0.17 | 11.16 |
|  | Chlorfenapyr | 2.05 | 1.80 to 2.34 | 2.82±0.25 | -1.86 |
|  | Pirimiphos-methyl | 1.05 | 0.90 to 1.21 | 3.03±0.26 | -1.62 |
| **Cyp9k1<sup>+</sup>** | Indoxacarb | 11.40 | 8.78 to 14.84 | 1.38±0.19 | 10.20 |
|  | Chlorfenapyr | 1.59 | 1.22 to 2.07 | 2.04±0.24 | -2.40 |
|  | Pirimiphos-methyl | 0.74 | 0.63 to 0.85 | 2.56±0.21 | -2.31 |

## Discussion

Mosquito population control tools that are based on insecticides have succeeded in preventing millions of malaria cases and deaths (2). However, the selection of high levels of insecticide resistance in *An. gambiae* populations, which is a result of the extensive use of these tools and our reliance on a few active ingredients, is now threatening their efficacy. Several studies have analyzed insecticide resistance in *An. gambiae* populations at a transcriptomic and genomic level, associating a plethora of genes and genomic alterations with the phenotype. However, very few of these genes and alterations, that are also frequently co-existing, have been functionally validated and it remains largely unknown what their effect size is.

We used the GAL4-UAS system to over-express in susceptible *An. gambiae* mosquitoes the *Cyp9k1* P450 and the *Abch2* transporter in susceptible *An. gambiae* mosquitoes. These two genes have been previously associated with resistance to pyrethroids, but without *in vivo* functional validation (12, 19, 32). *Cyp9k1* has been found over-expressed in several Anopheles resistant populations and the gene locus shows signals of positive selection (33, 34). *In vitro* metabolism assays have shown CYP9K1’s ability to metabolize deltamethrin and the insect growth regulator pyriproxyfen (12). In this study, we show that its over-expression in multiple tissues, under the polyubiquitin promoter results (29) in resistance to all pyrethroids tested at the WHO diagnostic dose, with the highest resistance ratio against *α*-cypermethrin (29.94-fold). A smaller, but significant increase in tolerance to DDT was also observed, while no resistance was detected to the organophosphates malathion and pirimiphos-methyl and the carbamate bendiocarb. The pyrethroid resistance profile of *Cyp9k1* over-expressing mosquitoes is similar to those previously generated in *Cyp6p3* and *Cyp6m2* over-expressing mosquitoes (all three-show resistance to permethrin and deltamethrin), but differs in relation to non-pyrethroid insecticides (14). *Cyp6p3* and *Cyp6m2* over-expressing mosquitoes did not show resistance to DDT and *Cyp6p3* mosquitoes, in contrast to *Cyp9k1*, showed resistance to bendiocarb. Thus, populations over-expressing both *Cyp6p3* and *Cyp9k1* could be multi-resistant to pyrethroids, DDT and bendiocarb.

We also functionally validated the role of the ABCH2 transporter. Transient silencing of the *Abch2* expression was previously shown to result in increased mortality upon deltamethrin exposure and increased penetration of the insecticide, as measured using radiolabeled insecticide (19). The gene was also found to be expressed in mosquitoes’ appendages and specifically in the epithelial cells of the legs, underneath the cuticle. Here we show that over-expression of *Abch2* in multiple tissues (including the appendages), under the polyubiquitin promoter did not confer resistance to the WHO diagnostic dose of any insecticide tested (19). However, we observed a clear reduction in deltamethrin sensitivity (resistance ratio 8-fold), when using three times lower deltamethrin dose. Thus, the over-expression of *Abch2* alone most likely confers only a mild insensitivity and may not be a primary resistance mechanism. However, we need to note that the use of the polyubiquitin promoter might not fully mirror the internal expression pattern (tissue and cell specific) of the genes tested and the levels of over-expression achieved might not be equivalent to those in resistant populations. Thus, the actual levels of resistance conferred by a gene, could be over or under estimated. In the future the system would greatly benefit from the availability of different promoters including tissue and cell specific, as well as inducible ones.

Resistant mosquitoes often harbor multiple resistance mechanisms (12, 22–25). This can arise through co-selection events, for example, when loci are physically close on the genome and belong to the same gene cluster exhibiting copy-number variation (35, 36), or when selection acts on a trans-regulator that alters the expression of several enzymes simultaneously (37–39). In such cases, functional redundancy may or may not be present. Multiple resistance mechanisms can also accumulate through sequential selection events. Here, selection pressure may come from exposure to different classes of insecticides, leading to the emergence of multi-resistant phenotypes. An example is the overexpression of detoxification enzymes that metabolize distinct insecticide classes, such as GSTE2, which metabolizes DDT (14, 16), and CYP6P3, which metabolizes pyrethroids and bendiocarb (14, 26). Another driver for the accumulation of resistance mechanisms would be the selection for increased resistance strength against a particular class or classes.

Here we demonstrate a synergistic interaction between target-site and metabolic resistance mechanisms. The combination of the L995F mutation in the VGSC with the over-expression of either *Cyp6p3* or *Cyp6m2* produced multiplicative resistance (RR_combined_ > RR_mechanism 1_ x RR_mechanism 2_). In the case of the Cyp6p3^+^+*kdr* combination we found significantly high resistance even to the 10× WHO diagnostic dose. This is in line with previous studies showing synergistic effects between target site and metabolic resistance mechanisms using *Drosophila* as a model organism (40) or in mosquitoes (10, 17, 41). In this study we expand this observation and show that combined over-expression of two detoxification enzymes (in this case of *Cyp6p3* with either *Abch2*, *Cyp9k1*, *Coeae6g* or the *Gste2*) can also result in significantly higher resistance levels than those conferred by each enzyme individually. In some combinations, the effect was particularly pronounced, such as the *Cyp6p3* and *Cyp9k1* combination, where resistance was observed not only to the 1× diagnostic dose of permethrin and deltamethrin, but also at the 5× diagnostic dose.

The mechanisms underlying these synergistic interactions remain poorly understood. For combinations of target-site mutations with over-expressed detoxification enzymes, one possibility is that the reduced binding affinity of the insecticide to the mutated VGSC allows more time for detoxification enzymes to metabolize the insecticide (if they are also over-expressed in the target tissues). Alternatively, detoxification enzymes may reduce the amount of insecticide that reaches the VGSCs in the target tissue, which combined with decreased binding affinity, results in a synergistic decrease in the channel’s inhibition. Similarly, how different detoxification enzymes act synergistically is not yet experimentally proven. In our system, the combined effect of P450s is unlikely to result simply from higher total P450 levels. This is supported by two observations: Firstly, Cyp6p3⁺+Cyp6p3⁺ mosquitoes, despite having higher *Cyp6p3* expression than either GAL4-Cyp6p3⁺ or Cyp6p3⁺ mosquitoes, were not more resistant, and secondly, Cyp6p3^+^+Cyp9k1^+^ mosquitoes were more resistant compared to Cyp6p3^+^, despite having lower *Cyp6p3* expression and similar *Cyp9k1* expression. A plausible hypothesis on how different detoxification enzymes could act in combination is that they form interactomes with enhanced metabolic activity. Indeed, there are indications from research in mammals that P450s can interact and their complexes may have altered substrate turnover efficiency (42, 43). Alternatively, different detoxification enzymes may act sequentially in a metabolic cascade that enhances insecticide clearance (44). In such a scenario, the metabolite produced by one enzyme, which can retain toxicity, could serve as the substrate for another enzyme that may even exhibit higher affinity for the metabolite than for the parental compound (45). Sequential metabolism could also prevent the loss of catalytic efficiency that occurs when a single enzyme is required to process both the parental insecticide and its metabolites, which can compete for the active site (11). This mechanism has been proposed for CYP6M2, which metabolizes deltamethrin and has also been shown to further transform some of its initial metabolites (11). Furthermore, is has been shown that while CYP6Z2, another P450 frequently overexpressed in pyrethroid resistant mosquitoes, cannot metabolise native pyrethroids, it metabolises PBAlc and PBAld, the major metabolites produced by other pyrethroid metabolizing P450s (46).

We also showed that when the over-expression of *Abch2* is combined with the over-expression of a metabolic enzyme, in this case CYP6P3, it can increase the levels of resistance conferred by the P450. In this combination the two mechanisms could act independently; the ABCH2 may export a portion of the insecticide not permitting its internalization, while CYP6P3 metabolizes the internalized molecules. Therefore, when combined they achieve a greater reduction in the number of active molecules reaching the nervous system.

We also assessed the sensitivity of Cyp6p3^+^ and Cyp9k1^+^ mosquitoes to the pro-insecticides chlorfenapyr, pirimiphos-methyl and indoxacarb. Mosquitoes over-expressing P450s are expected to be more vulnerable to pro-insecticides that require P450 mediated activation. Both CYP6P3 and CYP9K1 have previously been shown to activate the mitochondrial blocker chlorfenapyr *in vitro*, by generating the metabolite tralopyril (31). Our *in vivo* data build on these *in vitro* findings, to show that when *Cyp6p3* and *Cyp9k1* are overexpressed in mosquitoes there is an increase in chlorphenapyr sensitivity relative to controls. An increase in sensitivity was also observed to pirimiphos-methyl, despite the production of an oxidatively cleaved metabolite alongside the active pirimiphos-methyl *in vitro*. This suggests that *in vivo* the rate of PM activation by these P450s exceeds the rate of its oxidative cleavage, at least when exposed by topical application. The relatively low increase in sensitivity we observed in Cyp6p3^+^ and Cyp9k1^+^ mosquitoes indicates that the endogenous P450 activity is likely sufficient to activate the pro-insecticides and that additional P450 expression, at least in the tissues dictated by the ubiquitin promoter, does not alter the balance between activation and detoxification.

In contrast to PM and chlorfenapyr, Cyp6p3^+^ and Cyp9k1^+^ mosquitoes exhibited approximately 10-fold resistance to indoxacarb. This was expected for CYP6P3 given that only a hydroxylated metabolite was identified *in vitro*. In the case of CYP9K1, where both a hydroxylated and the active DCJW were identified, the toxicity test implies that the generation of the less toxic hydroxylated metabolite is the predominant pathway *in vivo*. In contrast to the combined effect which we observed between *Cyp6p3* and *Cyp9k1* over-expression in conferring resistance to pyrethroids, only minor effects were observed for the balance of activation/detoxification of the pro-insecticides tested. This difference in indoxacarb metabolism indicates that the combined effect of different P450 co-expression is insecticide specific and may involve the action of the different detoxification enzymes at different sites of the insecticide molecule, which is in line with a sequential metabolism model.

The data presented in this study not only advance our understanding on the mechanisms of insecticide resistance and how these translate to phenotypic resistance, but also have important implications for the design of efficient insecticide resistance management strategies. The demonstrated combined effects of different resistance mechanisms underscore the need to revise both the design and interpretation of molecular diagnostics used in field populations. Currently all molecular diagnostics evaluate resistance markers (mutations, over-expression levels) independently and the interpretation of the results is restricted to the presence and frequency of resistant alleles (or over-expression rates of detox genes) (25, 47, 48). Our findings show, however, that it is the interaction among multiple mechanisms that ultimately shapes resistance levels. This strongly supports the development of multi-loci diagnostics capable of predicting not only the presence of resistance, but also its magnitude in field mosquito populations. Furthermore, our results highlight the importance of evaluating, *in vivo*, the presence of cross or negative-cross resistance to pro-insecticides, mediated by the over-expression of P450s circulating in field populations. Such assessments must be conducted on a case-by-case basis to avoid failure of resistance-management strategies that rely on pro-insecticide activation (in the case of cross resistance) and at the same time leverage the power of these compounds to combat highly pyrethroid resistant populations, in the case of negative cross resistance. Finally, this work has contributed important synergism data to the considerable progress made over the past decades in understanding insecticide resistance and developing evidence-based management strategies, however our knowledge of resistance is still incomplete. Further research is needed to fully elucidate the dynamics of the different mechanisms that underpin insecticide resistance.

## Materials and Methods

### Mosquito rearing

All mosquitoes were reared at 26±2 °C and 70±10% relative humidity under a 12:12 h light–dark photoperiod cycle. Larvae were fed ground fish food (Tetramin Tropical Flakes; Tetra, Blacksburg, VA, USA), and adults were maintained with 10% sucrose solution *ad libitum*.

### Creation of UAS-Abch2 and UAS-Cyp9k1 responder lines

Generation of UAS-Abch2 and UAS-Cyp9k1 responder lines followed a previously described approach. Briefly, the coding sequence of Abch2 (AGAP0003680, 2283bp) and Cyp9k1 (AGAP000818, 1596bp) was amplified from the susceptible Kisumu strain using Phusion High-Fidelity DNA polymerase (New England Biolabs, Herts, UK) and primers listed in Supplementary Table 4. The coding sequence was sub-cloned into the pGEM-T Easy vector (Promega, Madison, USA) and then cloned into the pSL*attB:YFP:Gyp:UAS14i:Gyp:attB plasmid, downstream of the UAS as a NheI/XhoI (for *Abch2*) and EcoRI/NcoI (for *Cyp9k1*) restriction enzyme fragment. The structure of the responding plasmids is shown in Supplementary Figure 1A and 1B. Plasmids were sequenced to verify no differences in the cloned sequences compared to Vector base reference sequences.

To generate the responder lines, embryos from the A11 docking line, carrying a 3xP3-eCFP marker and 2 inverted attP sites, were injected with 150 ng/μL of the integrase helper plasmid pKC40 (encoding phiC31 integrase) mixed with 350 ng/μL of the responder plasmid, following previously described procedures (14, ^29^). Transient positive F_0_ individuals (YFP signal in their anal papillae) were sexed prior to emergence, pooled into founder cages and outcrossed with wild-type G3 mosquitoes. To obtain homozygous responder lines virgin F_1_ progeny carrying exclusively the YFP marker (indicative of cassette exchange) were crossed to males of the parental A11 docking line (labelled with CFP). F_2_ progeny were intercrossed, and homozygous UAS-Abch2 and UAS-Cyp9k1 strains were established by selecting individuals carrying only the YFP marker. Details of embryo injections and the generation of the UAS-Abch2 and UAS-Cyp9k1 responder lines are provided in Supplementary Figure 1A, 1B and Supplementary Table 2.

### Creation of the Ubi-A10: UAS Cyp6p3 line

To generate a line that natively over-expresses *Cyp6p3* under the Ubiquitin promoter the responder plasmid generated and used to make the UAS-Cyp6p3 line in Adolfi et al.,2019 was injected in embryos of the Ubi-A10 driver line(14), carrying a 3xP3-eCFP marker and 2 inverted attP sites, using the same procedures described above. Transiently positive F_0_ individuals were sexed as pupae and outcrossed with G3 mosquitoes. F_1_ progeny carrying both YFP and CFP markers in the eyes and nerve cord (indicative of cassette integration) were retained and outcrossed with G3 to expand the strain for one generation, while wild-type individuals lacking fluorescence were removed each generation. Details of embryo injections and the generation of Ubi-A10: UAS Cyp6p3 driver line are provided in Supplementary Table 2 and Supplementary Figure 4A.

### Establishment of Ubi-GAL4+kdr, UAS-Cyp6p3+kdr and UAS-Cyp6m2+kdr lines

The Kisumu^995F^ (*kdr*) line generated and described in Grigoraki et al.,2021 (10) was crossed with the Ubi-GAL4 driver line (14) (CFP^+^ marked) and the UAS-Cyp6p3 and UAS-Cyp6m2 responder lines (14) (YFP^+^ marked), respectively. Progeny of the crosses (F_1_) were backcrossed to the Kisumu^995F^ line to obtain F_2_ homozygotes for the mutation. F_2_ individuals were screened for the presence of the YFP^+^ or CFP^+^ markers. Individuals positive for the fluorescent signal were retained and left to emerge individually. The pupae case was used to extract DNA and select homozygotes for the *kdr* mutation using an LNA diagnostic assay, as previously described (10). Homozygotes of the *kdr* were intercrossed and thereafter kept as a mix of heterozygotes and homozygous for the GAL4 and UAS transgenic cassettes (Ubi-GAL4+*kdr*, UAS-Cyp6p3+*kdr* and UAS-Cyp6m2+*kdr*) (Supplementary figure 5A-C). Screening and selection of YFP^+^ and CFP^+^ positive individuals was performed each generation to enrich the frequency of the transgenic cassettes. To obtain individuals that over express the *Cyp6p3* or *Cyp6m2* and are homozygous for the *kdr* mutation, YFP^+^ individuals from the UAS-Cyp6p3+*kdr* and UAS-Cyp6m2+*kdr* lines were crossed to CFP^+^ individuals from the Ubi-GAL4+*kdr* line. Progenies that were YFP and CFP both positive were used in WHO toxicity bioassays (Supplementary figure 5D-E).

### Relative gene expression analysis

For gene expression analysis, total RNA was extracted from progeny of the different crosses using TRIzol reagent (Thermo Fisher Scientific, Waltham, USA) according to the manufacturer’s instructions, with at least three biological replicates of six 3∼5-day-old adults each. Genomic DNA was removed by treatment with RQ1 RNase-Free DNase (Promega). RNA concentration and purity were determined using a NanoDrop spectrophotometer, and first-strand cDNA was synthesized from 1 µg total RNA with the Engineered M-MLV Reverse Transcriptase Basic Kit (EnzyQuest, Heraklion, Greece) following the manufacturer’s instructions. RT-qPCR reactions were set up by using KAPA SYBR FAST (Merck, Rahway, USA) kit, RT–qPCR was performed on a Bio-Rad CFX96 (Bio-Rad Laboratories, Hercules, USA), as previously described (19), with three biological replicates and two technical replicates per biological replicate. The relative expression of each gene was analyzed by using the Pfaffl (49) method with the *Ribosomal Protein S7* (AGAP010592) as reference gene (19). Graphs and statistical analyses were performed using GraphPad Prism version 8.2.1 (GraphPad Software; https://www.graphpad.com/), with Welch’s *t*-test and one-way ANOVA (Tukey’s test) applied as appropriate.

### Assessment of insecticide susceptibility

#### WHO tubes bioassays

To assess insecticide susceptibility in *An. gambiae*, WHO tube bioassays were conducted following standard protocols (30) using insecticide-impregnated papers (Vector Control Research Unit, University Sains Malaysia), including 0.05%, 0.25% and 0.50% of deltamethrin, 0.75%, 3.75% and 7.50% of permethrin, 0.05% of *α*-cypermethrin, 0.1% of bendiocarb, 4% of DDT, 5% of malathion and 0.25% of pirimiphos-methyl, as previously described (10, 14). Different exposure times were used to estimate the LT_50_ of different insecticides, and PoLoPlus v2.0 (LeOra Software, Petaluma, USA) was used to calculate the LT_50_ value, Resistance Ratio and associated statistical parameters. For each time point, at least three replicates were performed, each consisting of approximately 20 females aged 3-5 days-old. The Knockdown rate was recorded immediately after the insecticide exposure and the mortality was recorded 24 h later. Welch’s *t* test was used to evaluate statistical differences between mortality rates. Statistics were calculated using a GraphPad Prism version 8.2.1 (GraphPad Software; https://www.graphpad.com/).

#### Topical application bioassays

For topical application bioassays, indoxacarb (98.32%), chlorfenapyr (99.38%), and pirimiphos-methyl (97.00%) were purchased from Dr. Ehrenstorfer (LGC Labo, Augsburg, DE) and dissolved in acetone to prepare stock solutions at 10,000 mg L⁻¹. Working solutions of the desired concentrations were subsequently prepared by serial dilution with acetone. Female mosquitoes (3-5 day-old) were anesthetized on ice for 2 min, and 0.2 µL of each insecticide solution (and acetone control) was applied to the surface of the pronotum using a PB600-1 repeating dispenser (Hamilton Company, Reno, USA). For each dose at least three replicates of 20 individuals each were tested. Mortality was assessed 24 h post-treatment, and bioassay data were analyzed using PoloPlus v2.0.

### Functional Expression of Cytochrome P450s

Cytochrome P450 (P450) gene sequences were synthesized into the pCW-OmpA2 expression vector, while *Anopheles gambiae* cytochrome P450 reductase (*Ag*CPR) was cloned into pACYC, as described by Mclaughlin et al., 2008 (50). Co-expression of each P450 enzyme with *Ag*CPR was performed in *E. coli* BL21 (DE3) Star cells. Membrane preparations were resuspended in 1× TSE buffer and stored at –80 °C. P450 content was quantified by CO-difference spectroscopy (51), and CPR activity was determined through NADPH-dependent cytochrome c reduction (52). Total protein concentration was measured using the Bradford assay (53).

To verify enzyme functionality, metabolism assays were conducted using permethrin and deltamethrin, which are known to be metabolized by CYP6P3 and CYP9K1 n (20 μM), respectively (26). Each reaction (100 ul final volume) contained 20 pmol recombinant CYP, 200 pmol An. gambiae cytochrome b5 (AgCytb5) (Stevenson et al., 2011 (11)), 20 μM insecticide and an NADPH-generating system consisting of 1 mM glucose-6-phosphate, 0.1 mM NADP+ and 1 unit/ml glucose-6-phosphate dehydrogenase (G6PDH) in 0.2 M Tris–HCl buffer (pH 7.4) with 0.25 mM MgCl₂. Incubations were carried out at 30 °C and 1250 rpm. Reactions were terminated by the addition of 100 ul acetonitrile, centrifuged at 13.000 rpm to remove precipitated proteins, and 100 ul from the resulting supernatants were analyzed by HPLC using a C18 reverse-phase column. Insecticides were eluted under isocratic conditions with water/acetonitrile mixtures (20:80 for permethrin and 10:90 for deltamethrin) at a flow rate of 1.25 mL/min for 20 minutes. Reactions were monitored by changes in absorbance between 225–250 nm and quantified by peak integration (Chromeleon, Dionex).

### Cytochrome P450 Metabolism Assays for Indoxacarb and Pirimiphos-methyl

Stock solutions of 10 mM indoxacarb (99,9 % purity; Sigma-Aldrich, USA) and pirimiphos-methyl (99,9 % purity; Sigma-Aldrich, USA) were prepared in acetonitrile.

a. Indoxacarb and CYP6P3: Reactions of 100 ul contained 10Μm indoxacarb final organic solvent concentration of 1 % (v/v)), 10 pmol CYP6P3 bacterial membrane protein, and 100 pmol AgCytb5 in 100 μL Tris–HCl buffer (0.2 M, pH 7.4) containing 0.25 mM MgCl₂.
b. Pirimiphos-methyl and CYP9K1: Reactions of 100 ul contained 25 μM pirimiphos-methyl (final organic solvent concentration of 2.5 % (v/v)), 25 pmol CYP9K1 bacterial membrane protein, and 250 pmol AgCytb5 in 100 μL Tris–HCl buffer (0.2 M, pH 7.4) containing 0.25 mM MgCl₂

Each incubation was performed in the presence and absence of an NADPH-generating system consisting of 1 mM glucose-6-phosphate, 0.1 mM NADP⁺, and 1 unit/mL glucose-6-phosphate dehydrogenase (G6PDH; Sigma-Aldrich, USA). Reactions were incubated at 30 °C with 1250 rpm oscillation, and stopped at 0 h and 2 h time points by adding 100 μL acetonitrile. Samples were mixed and stirred for an additional 30 min to ensure complete quenching. The mixtures were centrifuged at 10,000 rpm for 10 min, and the resulting supernatants were transferred to LC vials. An aliquot of 10 μL of each supernatant was injected for LC–MS analysis using an Ultimate 3000 Autosampler (Thermo Scientific, USA). All reactions were performed in triplicates, and results were compared with negative control reactions without the NADPH-regenerating system to assess substrate depletion.

### Identification of Indoxacarb and Pirimiphos-methyl metabolites by LC–MS analysis

The reactions of CYP6P3 and CYP9K1 with indoxacarb and pirimiphos-methyl, respectively, in the presence and absence of an NADPH-generating system, were analyzed using a high-resolution HPLC-MS/MS system. Sample injections of 10 μL were performed using an Ultimate 3000 Autosampler (Thermo Scientific, USA). Chromatographic separation was achieved on an Ultimate 3000 system (Thermo Scientific, USA) equipped with a 3 μm C18 (100×2.1 mm) reverse-phase analytical column (Fortis Technologies, UK).

a. Indoxacarb analysis: Indoxacarb reaction mixtures were analyzed with an isocratic mobile phase consisting of 70 % acetonitrile and 30 % H₂O, at a flow rate of 0.2 mL min⁻¹ over a 10-minute run. Detection was performed using an Electrospray Ionization (ESI) Q Exactive Hybrid Quadrupole-Orbitrap Mass Spectrometer (Thermo Scientific, USA) operated in positive ion mode. Mass spectrometric parameters were as follows: spray voltage 3500 V, sheath gas pressure 20 arbitrary units., auxiliary gas pressure 25 arbitrary units., and ion-transfer capillary temperature 350 °C. Parallel Reaction Monitoring (PRM) of indoxacarb was performed using collision-induced dissociation (CID) at 10–20 eV, with high-purity nitrogen as the sheath, auxiliary, and collision gas.
b. Pirimiphos-methyl analysis: Pirimiphos-methyl reaction mixtures were analyzed with an isocratic mobile phase consisting of 70% acetonitrile containing 0.1 % formic acid and 30 % H₂O containing 0.1 % formic acid, at a flow rate of 0.3 mL min⁻¹ over a 7-minute run. Detection was carried out using the same ESI Q Exactive Hybrid Quadrupole-Orbitrap MS system in positive ion mode. Instrument parameters were: spray voltage 4000 V, sheath gas pressure 40 arbitrary units., auxiliary gas pressure 5 arbitrary units., and ion-transfer capillary temperature 300 °C. PRM was conducted using CID at 10–20 eV under high-purity nitrogen.

The instrument was operated in full-scan, extracted-ion monitoring (EIM), and parallel reaction monitoring (PRM) modes. Data acquisition and processing were carried out using Xcalibur software (v4.0, Thermo Scientific, USA).

## Acknowledgments

We would like to thank Theodora Tatsiou for assisting with the generation of the UAS-Cyp9K1 plasmid used to generate the responder line and René Feyereisen for critically reviewing our manuscript. This publication is based on research funded by: the Hellenic Foundation for Research and Innovation (H.F.R.I.) under the: “3rd Call for H.F.R.I. Research Projects to support Post-Doctoral Researchers” (Project Number: 7406) (awarded to L.G) and the call “Basic research Financing (Horizontal support of all Sciences)” under the National Recovery and Resilience Plan “Greece 2.0” funded by the European Union – NextGenerationEU (H.F.R.I. Project Number: 16044) (awarded to J.V). Mengling Chen acknowledges the support of the China Scholarship Council program (Project ID: 202306990018), Latifa Remadi is supported by a Marie Sklodowska-Curie Postdoctoral Fellowship, Jocelyn M. F. Ooi and Mark J.I. Paine were funded by the IVCC (supported by the Bill and Melinda Gates Foundation under Grant Number OPP1148615) and the Medical Research Council (Grant Ref: MR/V001264/1).

