## Supplementary Information for "Synergistic action of different molecular mechanisms causes striking levels of insecticide resistance in the malaria vector *Anopheles gambiae*"

**Supplementary Table 1. Details of mosquitoes used in this study.**

| Mosquitoes used in this study | | | |
|---|---|---|---|
| **Line** | **Type** | **Fluorescent marker** | **Reference study** |
| A10 (Ubi-GAL4) | Driver line | $CFP^{+}$ | Adofi et al., 2018(1) |
| A11 | Docking line | $CFP^{+}$ | Adofi et al., 2018(1) |
| G3 | Susceptible strain | - | - |
| Kisumu | Susceptible strain | - | - |
| *Kdr* (Kisumu $^{995F}$) | CRISPR edited line with SNP | - | Grigoraki et al., 2021(2) |
| UAS-Abch2 | Responder line | $YFP^{+}$ | Generated process described in supplementary figure 1A |
| UAS-Cyp6p3 | Responder line | $YFP^{+}$ | Adofi et al., 2019(3) |
| UAS-Cyp9k1 | Responder line | $YFP^{+}$ | Generated process described in supplementary figure 1B |
| UAS-Gste2 | Responder line | $YFP^{+}$ | Adofi et al., 2019(3) |
| UAS-Coeae6g | Responder line | $YFP^{+}$ | Balaska et al., 2025(4) |
| Ubi-GAL4: UAS-Cyp6p3 | Driver line | $CFP^{+}$&$YFP^{+}$ | Generated process described in supplementary figure 4A |
| Ubi-GAL4+*kdr* | Driver line | $CFP^{+}$ | Generated process described in supplementary figure 5A |
| UAS-Cyp6p3+*kdr* | Responder line | $YFP^{+}$ | Generated process described in supplementary figure 5B |
| UAS-Cyp6m2+*kdr* | Responder line | $YFP^{+}$ | Generated process described in supplementary figure 5C |

| **GAL4/UAS progeny** | **Parental lines crossed** | **Genes being over-expressed** |
|---|---|---|
| $Abch2^{+}$ | Cross of Ubi-GAL4 and UAS-Abch2 | *Abch2* |
| $Cyp6p3^{+}$ | Cross of Ubi-GAL4 and UAS-Cyp6p3 | *Cyp6p3* |
| $Abch2^{+}$+$Cyp6p3^{+}$ | Cross of Ubi-GAL4: UAS-Cyp6p3 and UAS-Abch2 | *Abch2* and *Cyp6p3* |
| GAL4-$Cyp6p3^{+}$ | Cross of Ubi-GAL4: UAS-Cyp6p3 and G3 | *Cyp6p3* (single) |
| $Cyp6p3^{+}$+$Cyp6p3^{+}$ | Cross of Ubi-GAL4: UAS-Cyp6p3 and UAS-Cyp6p3 | *Cyp6p3* (double) |
| $Cyp9k1^{+}$ | Cross of Ubi-GAL4 and UAS-Cyp9k1 | *Cyp9k1* |
| $Cyp9k1^{+}$+$Cyp6p3^{+}$ | Cross of Ubi-GAL4: UAS-Cyp6p3 and UAS-Cyp9k1 | *Cyp9k1* and *Cyp6p3* |
| $Gste2^{+}$ | Cross of Ubi-GAL4 and UAS-Gste2 | *Gste2* |
| $Gste2^{+}$+$Cyp6p3^{+}$ | Cross of Ubi-GAL4: UAS-Cyp6p3 and UAS-Gste2 | *Gste2* and *Cyp6p3* |
| $Coeae6g^{+}$ | Cross of Ubi-GAL4 and UAS-Coeae6g | *Coeae6g* |
| $Coeae6g^{+}$+$Cyp6p3^{+}$ | Cross of Ubi-GAL4: UAS-Cyp6p3 and UAS-Coeae6g | *Coeae6g* and *Cyp6p3* |
| $Cyp6p3^{+}$+*kdr* | Cross of Ubi-GAL4+*kdr* and UAS-Cyp6p3+*kdr* | *Cyp6p3* with *kdr* mutation homozygosity |
| $Cyp6m2^{+}$+*kdr* | Cross of Ubi-GAL4+*kdr* and UAS-Cyp6m2+*kdr* | *Cyp6m2* with *kdr* mutation homozygosity |

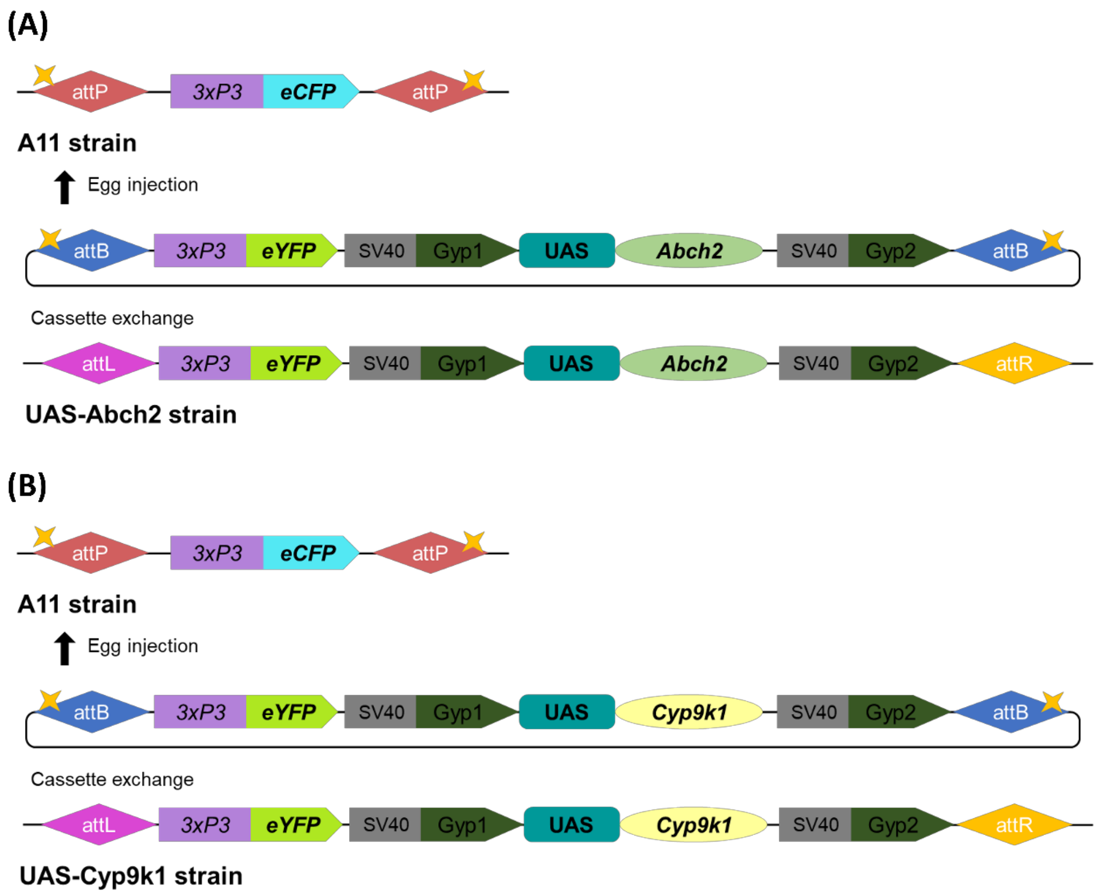


**Supplementary Figure 1. Schematic diagram showing the generation of UAS-Abch2 (A) and UAS-Cyp9k1(B) responder lines.**

**Supplementary Table 2. Summary of embryo injections, screening and crosses performed to generate the UAS-Abch2, UAS-Cyp9k1 and Ubi-GAL4: UAS-Cyp6p3 lines by RMCE.**

<table>
<tr><th colspan="6">UAS-ABCH2/ UAS-Cyp9k1/ Ubi-GAL4: UAS-Cyp6p3 injection data</th></tr>
<tr><td rowspan="6">UAS-Abch2 injection data</td><td>Total injected eggs on A11</td><td>Hatched Positive G<sub>0</sub> larvae</td><td>Hatched Negative G<sub>0</sub> larvae</td><td>Hatched rate</td><td>Positive rate</td></tr>
<tr><td>602</td><td>54</td><td>60</td><td>18.94%</td><td>47.37%</td></tr>
<tr><td colspan="2">Positive G<sub>0</sub> (only YFP<sup>+</sup>) Back-Cross with G3</td><td colspan="3">Test of F<sub>1</sub></td></tr>
<tr><td>Abch2♀: 30 X G3♂: 90</td><td>Abch2♂: 20 X G3♀: 60</td><td colspan="3">7 of F<sub>1</sub> progeny were identified with cassette exchange (YFP<sup>+</sup> only)</td></tr>
<tr><td rowspan="4">UAS-Cyp9k1 injection data</td><td>Total injected eggs on A11</td><td>Hatched Positive G<sub>0</sub> larvae</td><td>Hatched Negative G<sub>0</sub> larvae</td><td>Hatched rate</td><td>Positive rate</td></tr>
<tr><td>287</td><td>45</td><td>30</td><td>26.13%</td><td>60.00%</td></tr>
<tr><td colspan="2">Positive G<sub>0</sub> (only YFP<sup>+</sup>) Back-Cross with G3</td><td colspan="3">Test of F<sub>1</sub></td></tr>
<tr><td>Cyp9k1♀: 17 X G3♂: 51</td><td>Cyp9k1♂: 10 X G3♀: 30</td><td colspan="3">2 of F<sub>1</sub> progeny were identified with cassette exchange (YFP<sup>+</sup> only)</td></tr>
<tr><td rowspan="4">Ubi-GAL4: UAS-Cyp6p3 injection data</td><td>Total injected eggs on A10</td><td>Hatched Positive G<sub>0</sub> larvae</td><td>Hatched Negative G<sub>0</sub> larvae</td><td>Hatched rate</td><td>Positive rate</td></tr>
<tr><td>1600</td><td>220</td><td>N/A</td><td>N/A</td><td>N/A</td></tr>
<tr><td colspan="2">Positive G<sub>0</sub> (YFP<sup>+</sup> and CFP<sup>+</sup>) Back-Cross with G3</td><td colspan="3">Test of F<sub>1</sub></td></tr>
<tr><td>GAL4: Cyp6p3♀: 99 X G3♂: 273</td><td>GAL4: Cyp6p3♂: 120 X G3♀: 330</td><td colspan="3">16 of F<sub>1</sub> progeny were identified with cassette insertion (YFP<sup>+</sup> and CFP<sup>+</sup>)</td></tr>
</table>

N/A means values not recorded.

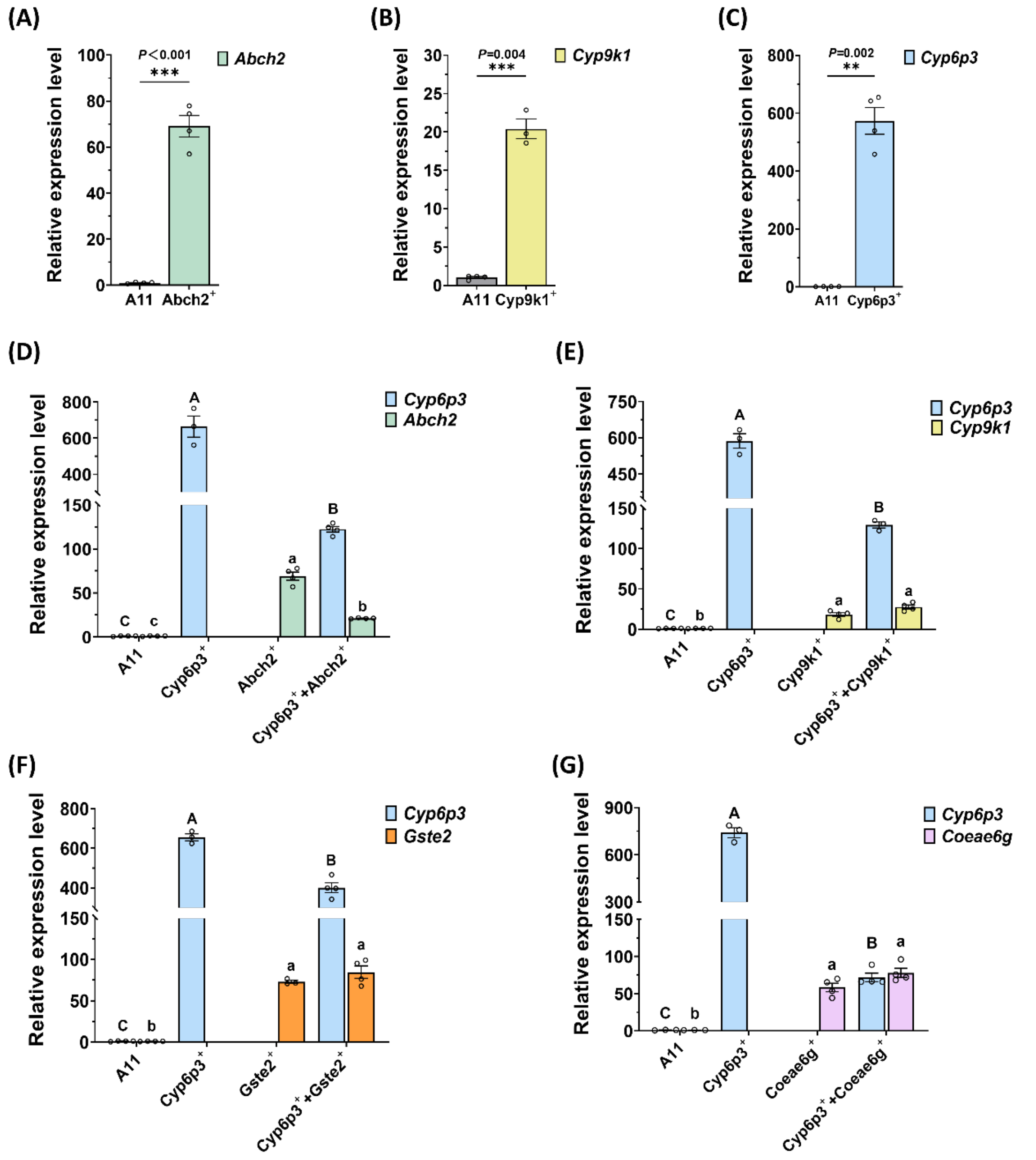


**Supplementary Figure 2. Relative expression levels of detoxification genes in GAL4/UAS progeny.**

Expression levels of (A) *Abch2* (Abch2$^{+}$), (B) *Cyp9k1* (Cyp9k1$^{+}$), (C) *Cyp6p3*(Cyp6p3$^{+}$) are provided relative to control A11 mosquitoes. Expression levels were analyzed using the Pfaffl method and the housekeeping *Ribosomal Protein S7* (AGAP010592) was used for normalization. Statistical significance was assessed using Welch's *t*-test (n.s., not significant, **P*< 0.05, ***P* < 0.01; ****P* < 0.001). (D) Expression level of *Cyp6p3* and *Abch2* in Cyp6p3$^{+}$/ Abch2$^{+}$ and Cyp6p3$^{+}$+Abch2$^{+}$ mosquitoes relative to A11 control strain. (E) Expression level of *Cyp6p3* and *Cyp9k1* in Cyp6p3$^{+}$/ Cyp9k1$^{+}$ and Cyp6p3$^{+}$+Cyp9k1$^{+}$ mosquitoes relative to A11 control strain. (F) Expression level of *Cyp6p3* and *Gste2* in Cyp6p3$^{+}$, Gste2$^{+}$ and Cyp6p3$^{+}$+Gste2$^{+}$ mosquitoes relative to A11 control strain. (G) Expression level of *Cyp6p3* and *Coeae6g* in Cyp6p3$^{+}$, Coeae6g$^{+}$ and Cyp6p3$^{+}$+Coeae6g$^{+}$ mosquitoes relative to A11 control strain. The *ribosomal protein S7* (AGAP010592) was used for normalization. Statistical significance was assessed using one-way ANOVA followed by a Tukey's test.

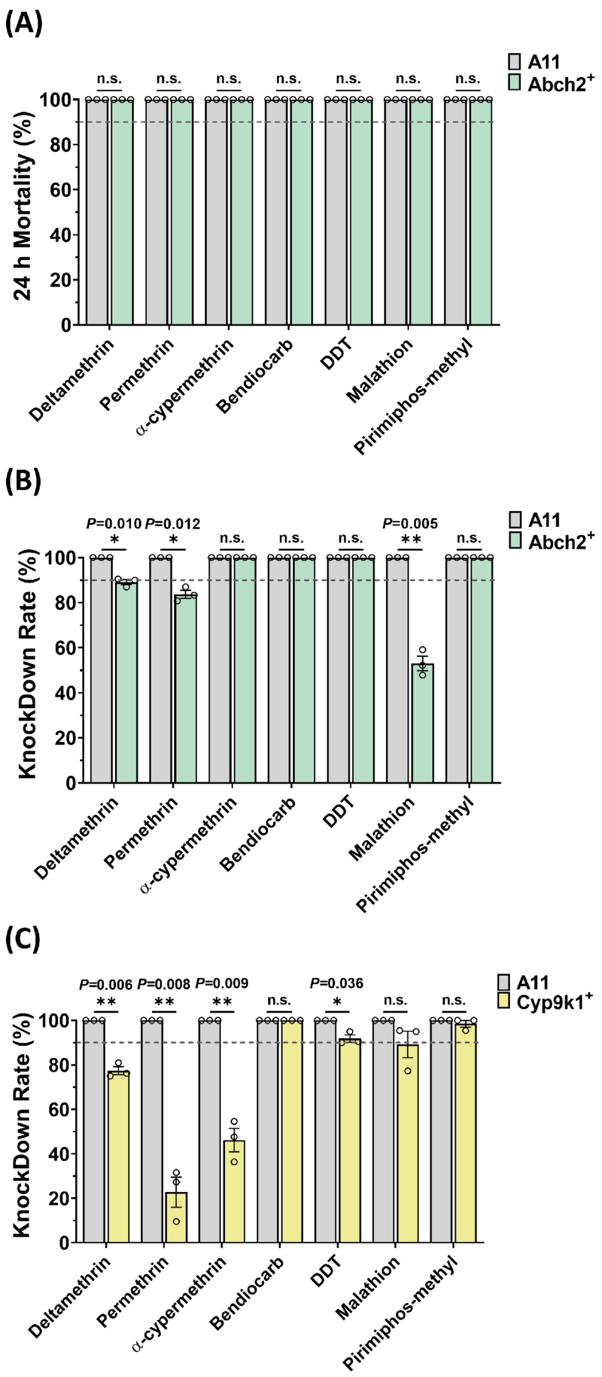


**Supplementary Figure 3. WHO toxicity bioassays for testing the effect of multi-tissue *Abch2* or *Cyp9k1* over-expression on insecticide resistance.**

(A) Percentage mortality of *Abch2* over-expressing (Abch2$^{+}$) and control (A11) mosquitoes 24 h after 1h exposure to the 1× WHO diagnostic dose of different insecticides (0.05% deltamethrin, 0.75% permethrin, 0.05% *α*-cypermethrin, 0.1% bendiocarb, 4% DDT, 5% malathion, and 0.25% pirimiphos-methyl). (B) The knockdown rate of *Abch2* over-expressing (Abch2$^{+}$) and control (A11) mosquitoes after the treatment of 1× WHO diagnostic dose of different insecticides. (C) The knockdown rate of *Cyp9k1* over-expressing (Cyp9k1$^{+}$) and control (A11) mosquitoes after the treatment of 1× WHO diagnostic dose of different insecticides. For all bioassays, at least three biological replicates were performed, each comprising 20–25 randomly selected 3~5-day-old females. Individual data points represent different biological replicates, and bars indicate the standard error (SE). The dotted line indicates the WHO 90% mortality threshold used to define resistance. Statistical significance was assessed using Welch's *t*-test (n.s., not significant, *$P$< 0.05, **$P$ < 0.01; ***$P$ < 0.001).

**Supplementary Table 3. Time-response toxicity assays for estimating resistance strength to 0.016% deltamethrin for *Abch2* over-expressing mosquitoes.**

The $LT_{50}$ (time required to obtain 50% mortality) is provided for each strain with the 95% fiducial limits (95% FL) and used for calculating the resistance ratio (RR) ($LT_{50}$ of Abch2$^{+}$ / $LT_{50}$ of A11). Each bioassay included at least five exposure time points, for each of which at three replicates, of 20–25, 3~5-day-old female mosquitoes were tested.

| Line | Insecticides | $LT_{50}$ | 95%FL | Slope±SE | RR |
|---|---|---|---|---|---|
| **A11** | Deltamethrin-0.016% | ~3 | | | |
| **Abch2$^{+}$** | Deltamethrin-0.016% | 25.07 | 21.67 to 29.24 | 2.17±0.18 | **~8.36** |

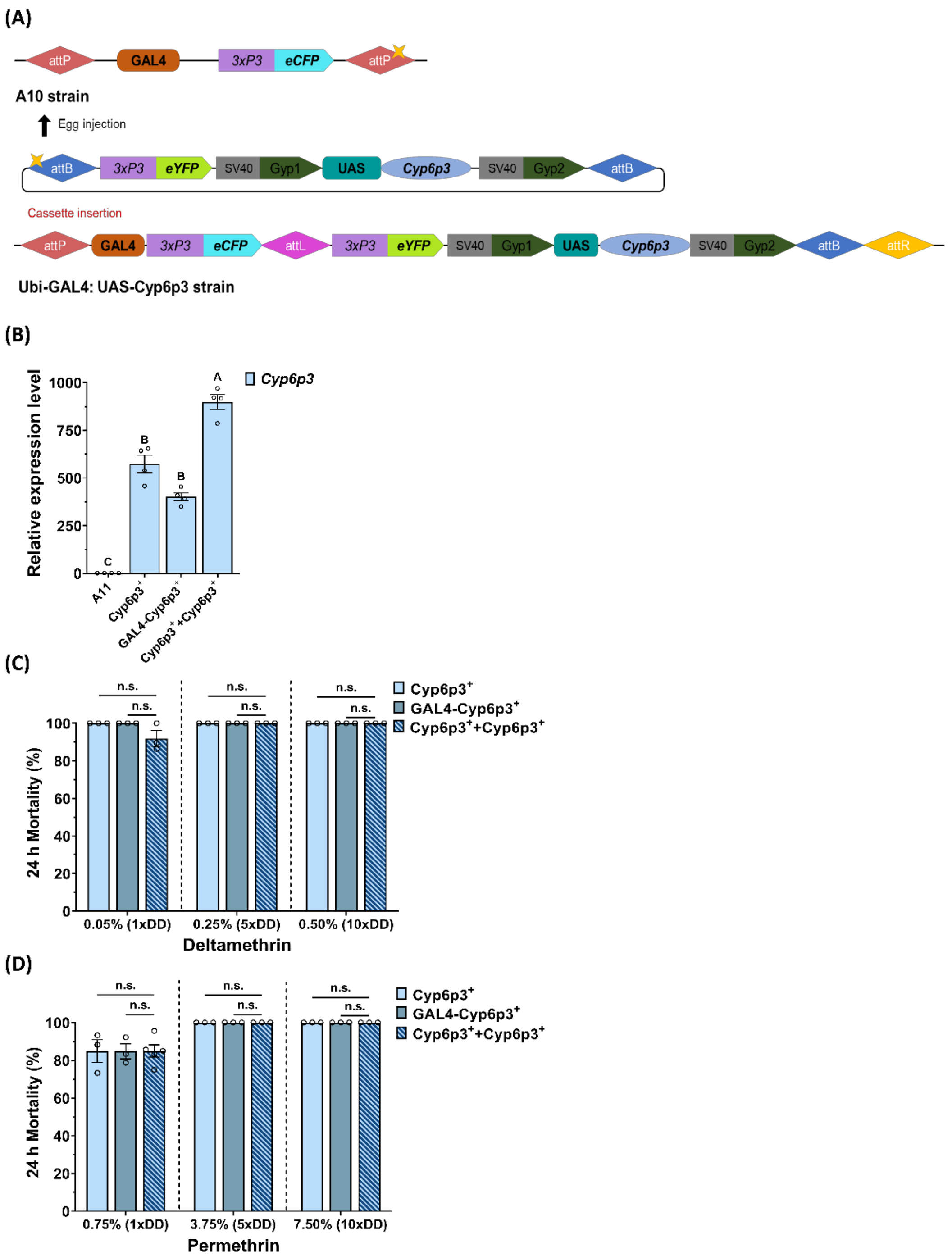


**Supplementary Figure 4. Generation of Ubi-GAL4: UAS-Cyp6p3 line and bioassay data.**

(A) Schematic diagram showing the generation of the Ubi-GAL4: UAS-Cyp6p3 line. (B) Expression level of Cyp6p3 in Cyp6p3$^+$(progeny of Ubi-Gal4 and UAS-Cyp6p3); GAL4-Cyp6p3$^+$ (progeny of Ubi-GAL4: UAS-Cyp6p3 with G3, harboring a single copy of the Cyp*6p3* transgene) and Cyp6p3$^+$+Cyp6p3$^+$ (progeny of the Ubi-GAL4: UAS-Cyp6p3 and UAS-Cyp6p3) mosquitoes relative to control A11. The *ribosomal protein S7* (AGAP010592) was used for normalization. Statistical significance was assessed using one-way ANOVA followed by a Tukey's test. (C) Percentage mortality of Cyp6p3$^+$, GAL4-Cyp6p3$^+$ and Cyp6p3$^+$+Cyp6p3$^+$ mosquitoes 24 h after 1h exposure to the 1× (0.05%), 5× (0.25%) and 10× (0.50%) WHO diagnostic dose of deltamethrin and (D) 1x (0.75%), 5x (3.75%) and 10x (7.5%) diagnostic dose of permethrin. For all bioassays, at least three replicates were performed, each comprising 20–25 3~5-day-old females. Individual data points represent different replicates, and bars indicate the standard error (SE). Statistical significance was assessed using Welch's *t*-test (n.s., not significant, *$P$< 0.05, **$P$ < 0.01; ***$P$ < 0.001).

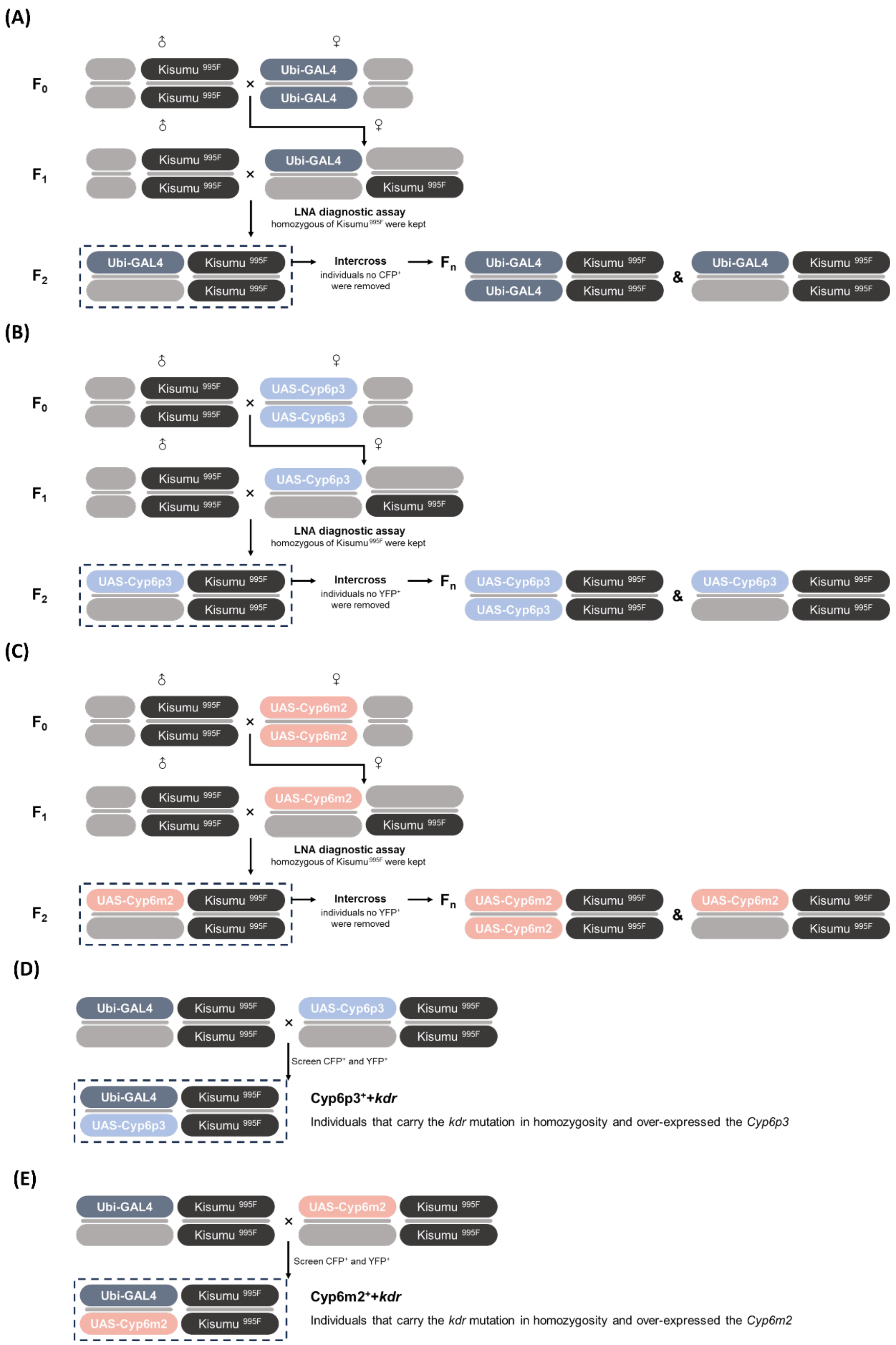


**Supplementary Figure 5. Schematic diagram showing the generation of Ubi-GAL4+*kdr* (A), UAS-Cyp6m2+*kdr* (B) and UAS-Cyp6p3+*kdr* (C) lines. Schematic diagrams showing the crosses that were performed to obtain individuals the carry the VGSC 995F mutation in homozygosity and over-express *Cyp6p3* (D) or *Cyp6m2* (E).**

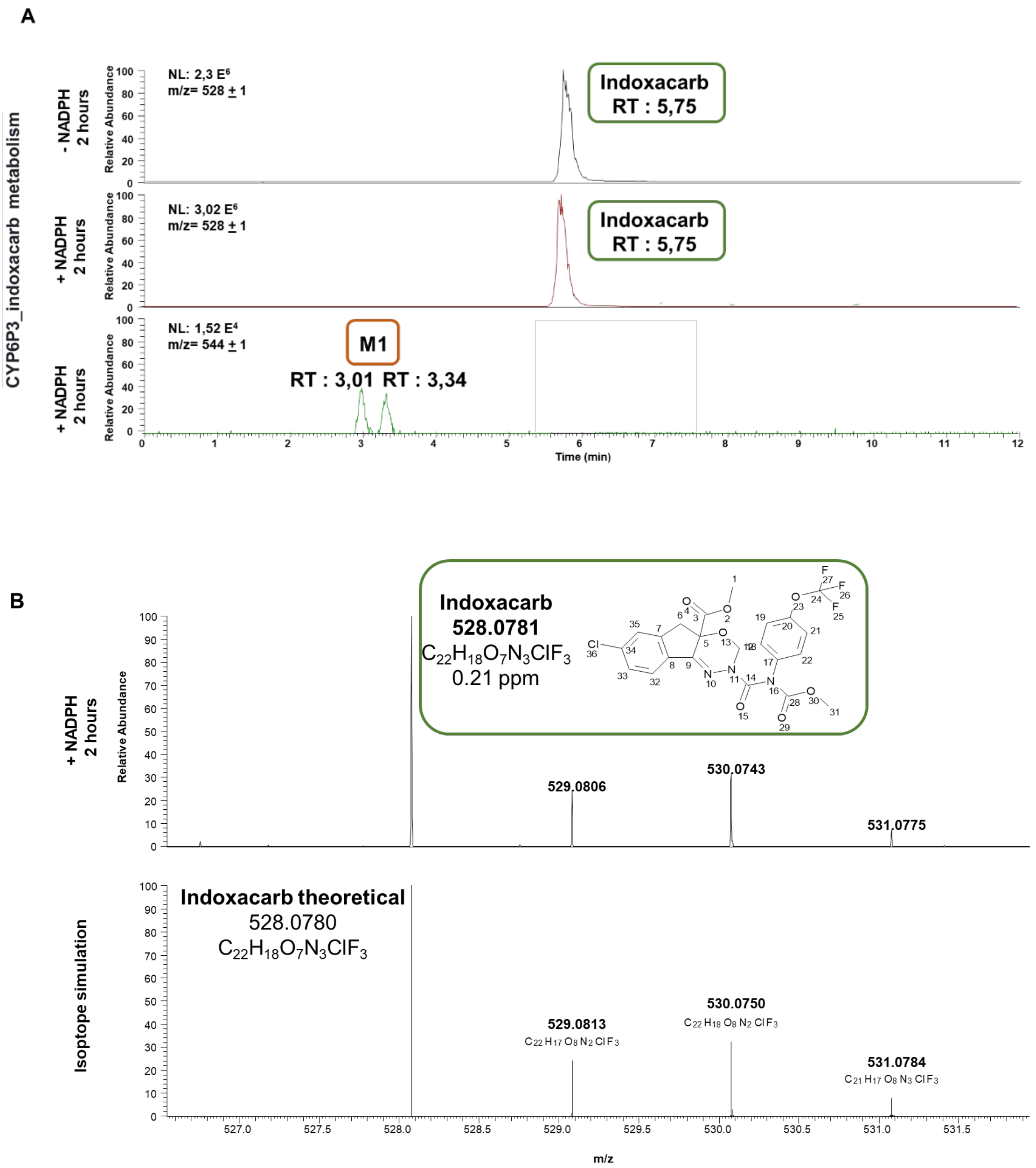
A
CYP6P3_indoxacarb metabolism
- NADPH 2 hours
NL: 2,3 E6
m/z= 528 ± 1
Indoxacarb
RT : 5,75
+ NADPH 2 hours
NL: 3,02 E6
m/z= 528 ± 1
Indoxacarb
RT : 5,75
+ NADPH 2 hours
NL: 1,52 E4
m/z= 544 ± 1
M1
RT : 3,01 RT : 3,34
Relative Abundance
Time (min)
B
+ NADPH 2 hours
Indoxacarb
528.0781
C22H18O7N3ClF3
0.21 ppm
529.0806
530.0743
531.0775
Isotope simulation
Indoxacarb theoretical
528.0780
C22H18O7N3ClF3
529.0813
C22 H17 O8 N2 Cl F3
530.0750
C22 H18 O8 N2 Cl F3
531.0784
C21 H17 O8 N3 Cl F3
m/z

C

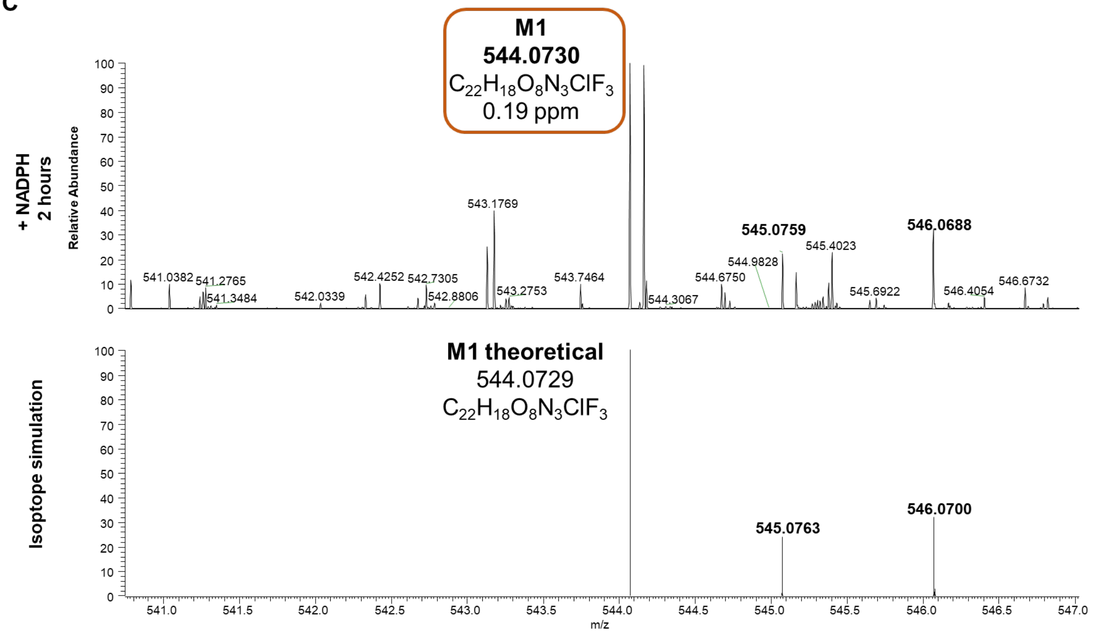


D

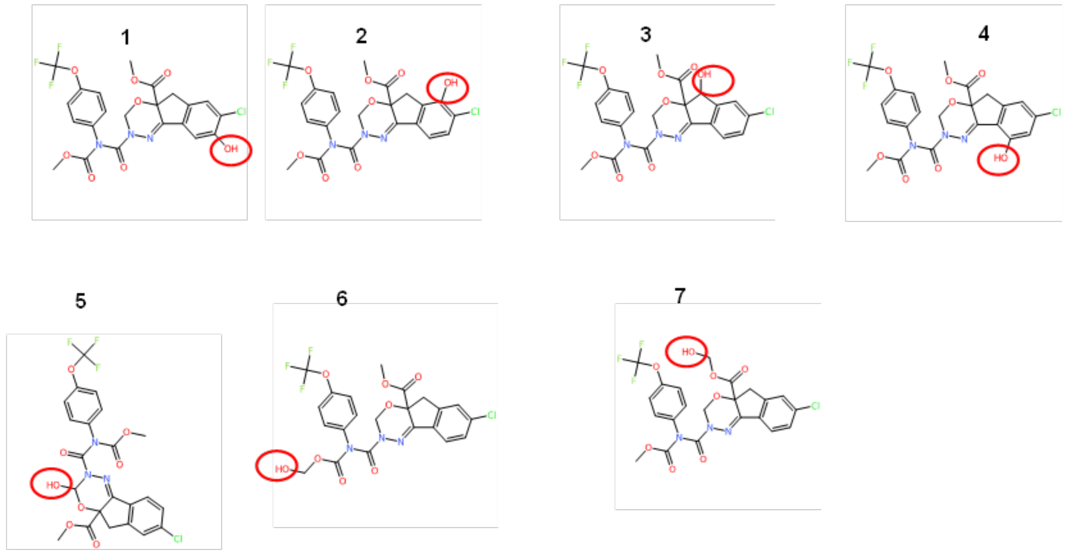


**Supplementary Figure 6. Metabolism of indoxacarb by recombinant CYP6P3.**

**(A)** LC extracted-ion chromatograms obtained using high-resolution full-scan MS showing an NADPH-dependent transformation of indoxacarb (eluting at 5.75 min) into a metabolite M1 (eluting at 3.01 and 3.34 min). **(B)** Accurate mass spectra of indoxacarb obtained using HPLC–electrospray ionization (ESI). Upper panel: high-resolution accurate mass of protonated indoxacarb at m/z $[M + H]^+ = 528.0781$. Lower panel: theoretical mass and isotope simulation of protonated indoxacarb at m/z $[M + H]^+ = 528.0780$. **(C)** Accurate mass spectra of metabolite M1 obtained using LC–ESI. Upper panel: high-resolution accurate mass of protonated indoxacarb metabolite M1 at m/z $[M + H]^+ = 544.0730$. Lower panel: theoretical mass and isotope simulation of the protonated metabolite at m/z $[M + H]^+ = 544.0729$. **(D)** Predicted structures of M1, the hydroxylated metabolite of indoxacarb, showing several possible hydroxylation sites as proposed by the Bio Transformer 3.0 tool. Accurate mass data, including elemental composition, theoretical and experimental m/z values, and mass accuracy (ppm), are reported for all compounds and metabolites to support structural assignments.

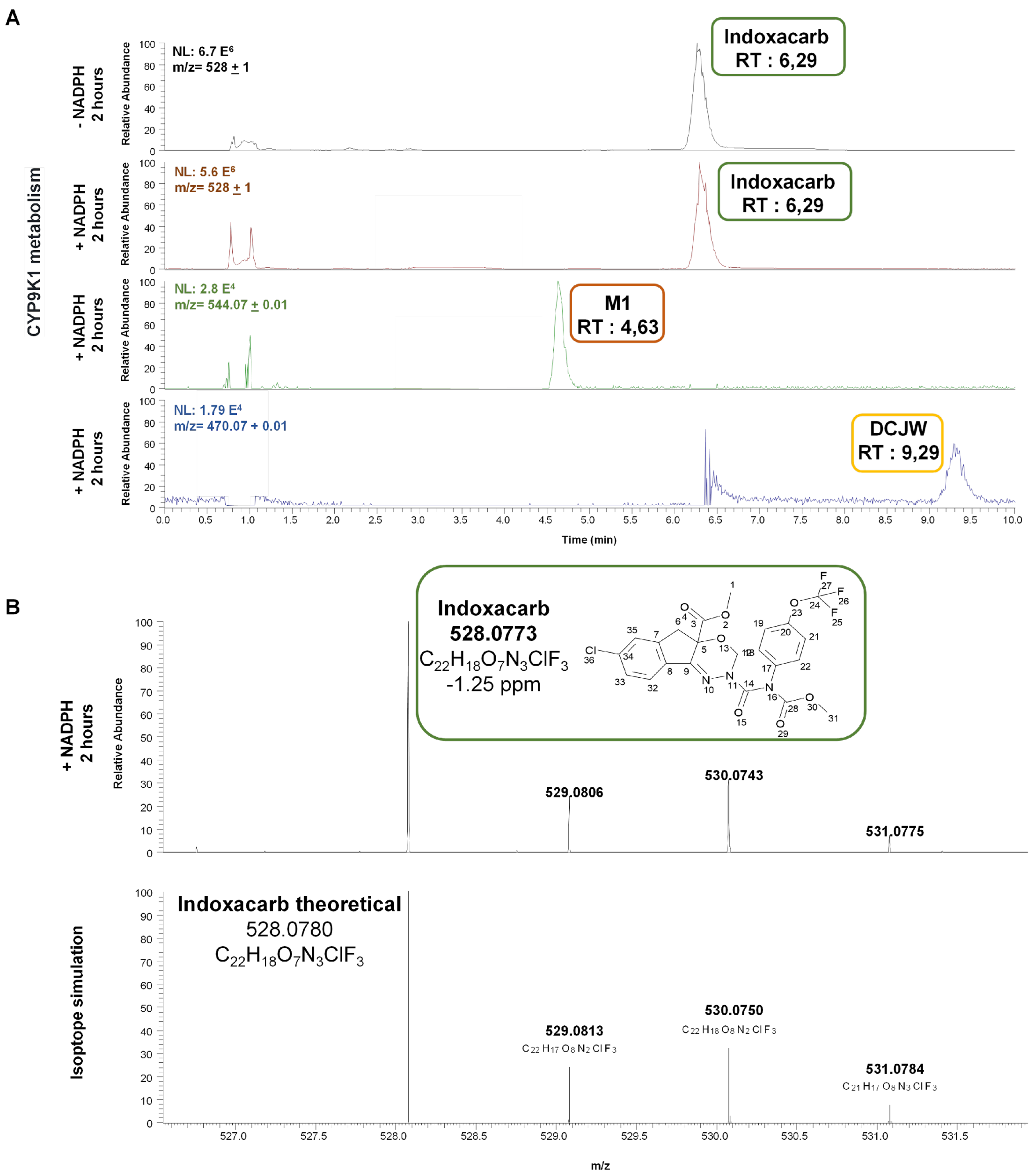
A
CYP9K1 metabolism
- NADPH 2 hours
NL: 6.7 E6
m/z= 528 ± 1
Indoxacarb RT : 6,29
+ NADPH 2 hours
NL: 5.6 E6
m/z= 528 ± 1
Indoxacarb RT : 6,29
NL: 2.8 E4
m/z= 544.07 ± 0.01
M1 RT : 4,63
NL: 1.79 E4
m/z= 470.07 + 0.01
DCJW RT : 9,29
Relative Abundance
Time (min)
B
+ NADPH 2 hours
Indoxacarb
528.0773
C22H18O7N3ClF3
-1.25 ppm
529.0806
530.0743
531.0775
Isoptope simulation
Indoxacarb theoretical
528.0780
C22H18O7N3ClF3
529.0813
C22 H17 O8 N2 Cl F3
530.0750
C22 H18 O8 N2 Cl F3
531.0784
C21 H17 O8 N3 Cl F3
m/z

C

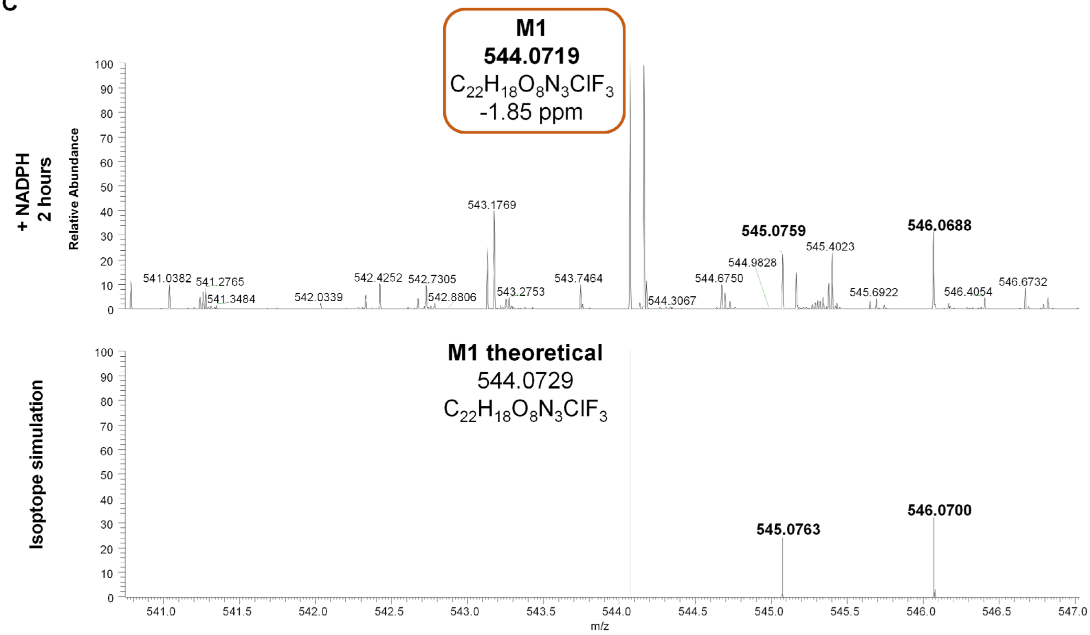
M1
544.0719
$C_{22}H_{18}O_8N_3ClF_3$
-1.85 ppm
+ NADPH
2 hours
Relative Abundance
543.1769
545.0759
546.0688
545.4023
544.9828
544.6750
541.0382
541.2765
541.3484
542.0339
542.4252
542.7305
542.8806
543.2753
543.7464
544.3067
545.6922
546.4054
546.6732
Isotope simulation
M1 theoretical
544.0729
$C_{22}H_{18}O_8N_3ClF_3$
545.0763
546.0700
m/z

D

1

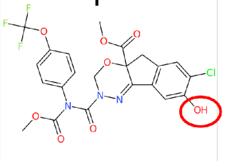

2

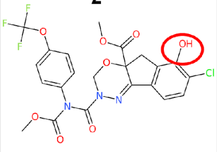

3

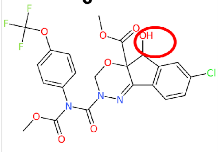

4

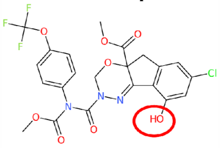

5

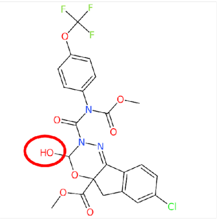

6

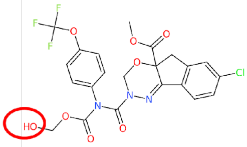

7

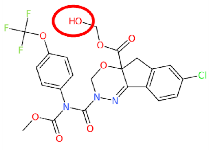

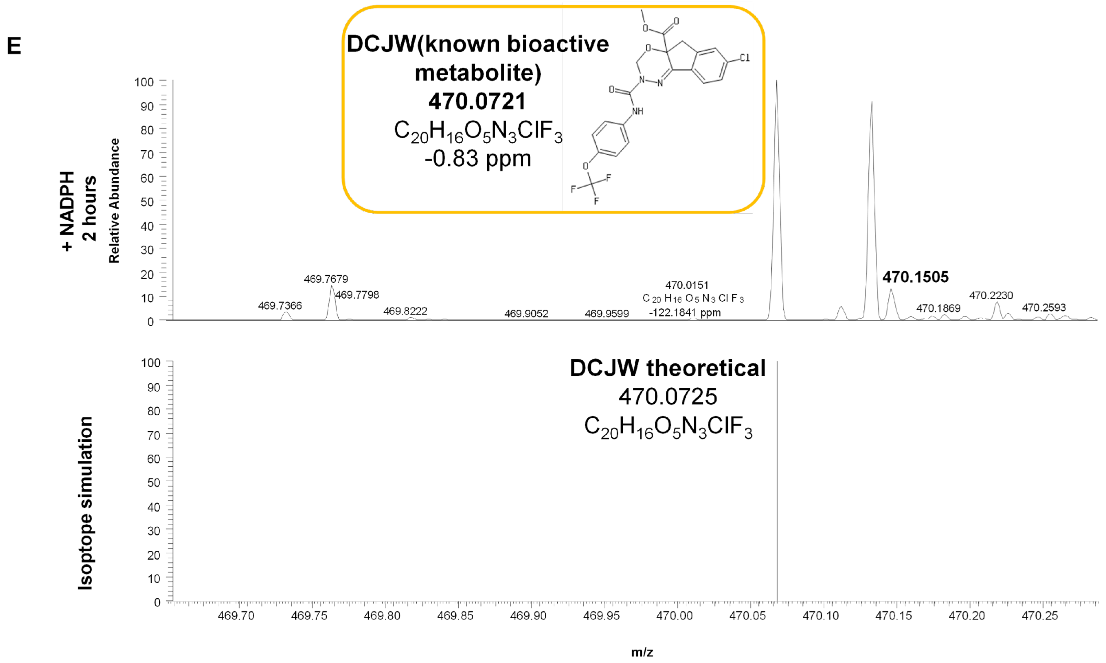


**Supplementary Figure 7**. **Metabolism of indoxacarb by recombinant CYP9K1.**

**(A)** LC extracted-ion chromatograms obtained using high-resolution full-scan MS showing an NADPH-dependent transformation of indoxacarb (eluting at 6.29 min) into the formation of an unidentified metabolite, M1 (eluting at 4.63 min), and the known bioactive N-decarbomethoxylated metabolite, DCJW (eluting at 9.29 min), when compared to the control reaction without NADPH. **(B)** Accurate mass spectra of indoxacarb obtained using HPLC–electrospray ionization (ESI). Upper panel: high-resolution accurate mass of protonated indoxacarb at m/z $[M + H]^+ = 528.0773$. Lower panel: theoretical mass and isotope simulation of protonated indoxacarb at m/z $[M + H]^+ = 528.0780$. **(C)** Accurate mass spectra of metabolite M1 obtained using LC–ESI. Upper panel: high-resolution accurate mass of protonated indoxacarb metabolite M1 at m/z $[M + H]^+ = 544.0719$. Lower panel: theoretical mass and isotope simulation of the protonated metabolite at m/z $[M + H]^+ = 544.0729$. **(D)** Predicted structures of M1, the hydroxylated metabolite of indoxacarb, showing several possible hydroxylation sites as proposed by the Bio Transformer 3.0 tool. Accurate mass data, including elemental composition, theoretical and experimental m/z values, and mass accuracy (ppm), are reported for all compounds and metabolites to support structural assignments. (**E**) The mass spectrum of the DCJW metabolite in positive ion mode showed a molecular ion peak at m/z $[M+H]^+ = 470.0721$, corresponding to a decarbomethoxylated product, 58 Da lower than the parent indoxacarb at m/z $[M+H]^+ = 528.0773$.

(A)

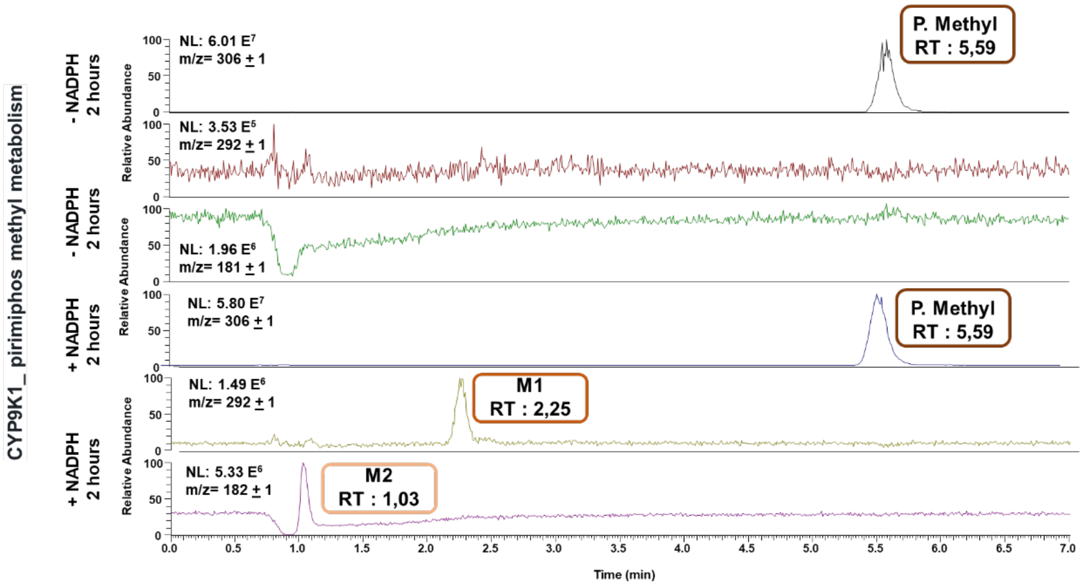

CYP9K1_ pirimiphos methyl metabolism
- NADPH 2 hours
+ NADPH 2 hours
Relative Abundance
NL: 6.01 E7
m/z= 306 ± 1
P. Methyl
RT : 5,59
NL: 3.53 E5
m/z= 292 ± 1
NL: 1.96 E6
m/z= 181 ± 1
NL: 5.80 E7
m/z= 306 ± 1
P. Methyl
RT : 5,59
NL: 1.49 E6
m/z= 292 ± 1
M1
RT : 2,25
NL: 5.33 E6
m/z= 182 ± 1
M2
RT : 1,03
Time (min)


(B)

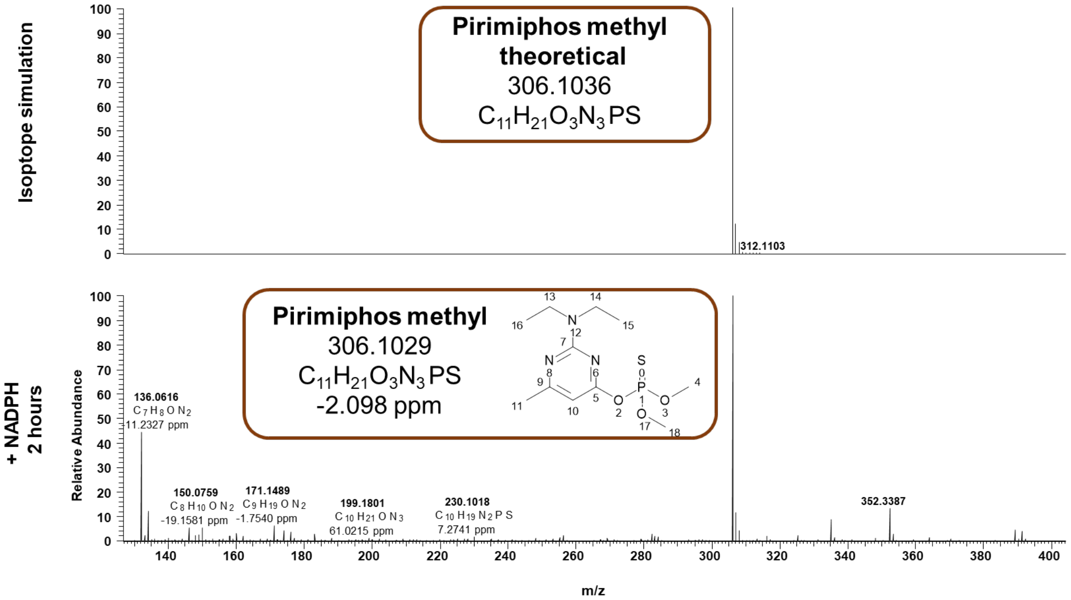

Isoptope simulation
Pirimiphos methyl theoretical
306.1036
C11H21O3N3PS
312.1103
+ NADPH 2 hours
Relative Abundance
Pirimiphos methyl
306.1029
C11H21O3N3PS
-2.098 ppm
136.0616
C7H8ON2
11.2327 ppm
150.0759
C8H10ON2
-19.1581 ppm
171.1489
C9H19ON2
-1.7540 ppm
199.1801
C10H21ON3
61.0215 ppm
230.1018
C10H19N2PS
7.2741 ppm
352.3387
m/z

(C)

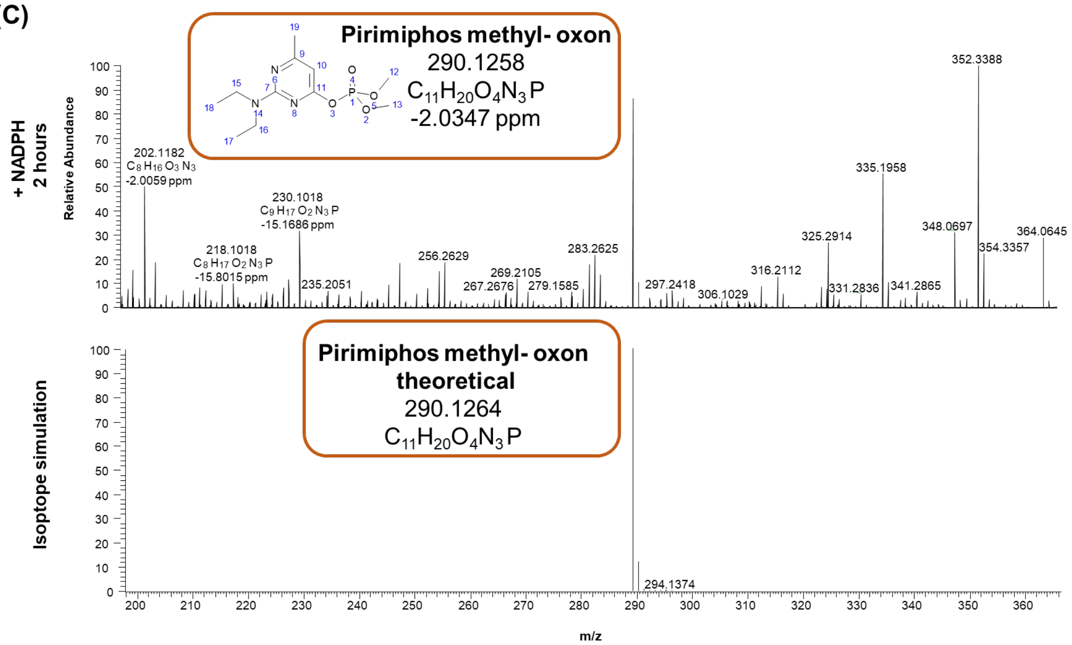


(D)

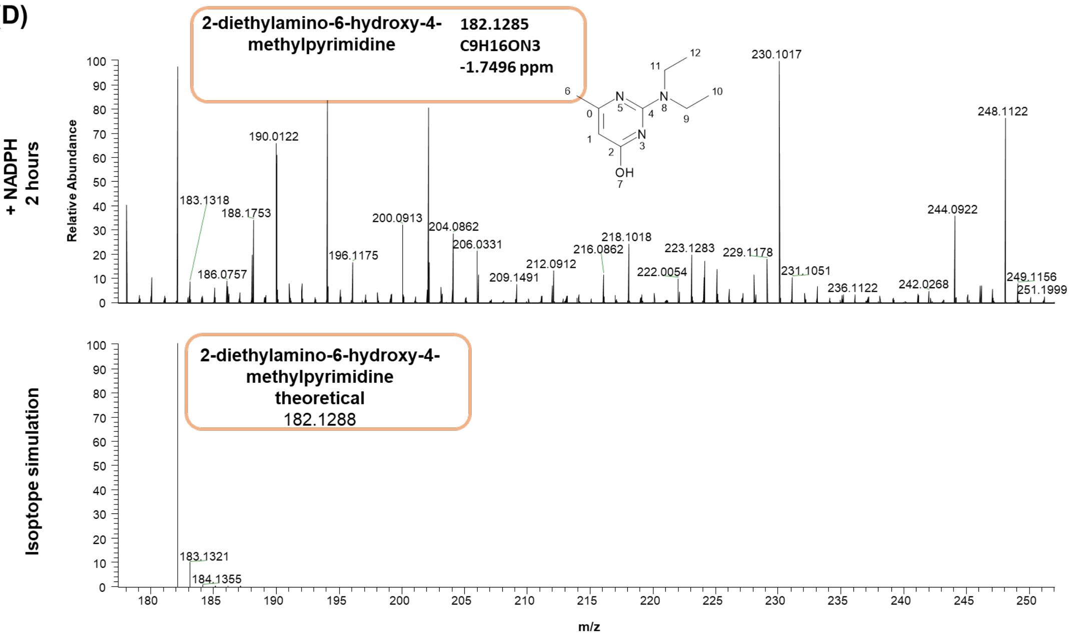


**Supplementary Figure 8. Metabolism of pirimiphos-methyl by recombinant CYP9K1.**

**(A)** LC extracted-ion chromatograms obtained using high-resolution full-scan MS showing an NADPH-dependent transformation of pirimiphos-methyl (P. methyl) by recombinant CYP9K1 expressed in *E. coli* membranes. After 2h of incubation, a 35.9 % depletion of the parent compound (eluting at 5.58 min) was observed, accompanied by the formation of two metabolites, M1 (2.25 min) and M2 (1.03 min), compared with the control reaction lacking NADPH. **(B)** Accurate mass spectra of parent compound, pirimiphos-methyl, obtained using HPLC–electrospray ionization (ESI). Upper panel: theoretical mass and isotope simulation of protonated pirimiphos methyl at m/z $[M + H]^+ = 306.1036$, Lower panel: high-resolution accurate mass of protonated pirimiphos methyl at m/z $[M + H]^+ = 306.1029$. **(C)** Accurate mass spectra of metabolite M1 obtained using LC–ESI. Upper panel: high-resolution accurate mass of protonated pirimiphos methyl metabolite M1 at m/z $[M + H]^+ = 290.1258$, Lower panel: theoretical mass and isotope simulation of the protonated pirimiphos methyl oxon at m/z $[M + H]^+ = 290.1264$. Metabolite M1 displayed a peak at m/z $[M + H]^+ = 290.1258$, which is 15.98 Da lower than the parent compound, consistent with a desulfurization (S → O) reaction forming pirimiphos-methyl oxon. **(D)** Accurate mass spectra of metabolite M2 obtained using LC–ESI. Upper panel: high-resolution accurate mass of protonated pirimiphos methyl metabolite M2 at m/z $[M + H]^+ = 182.1285$, Lower panel: theoretical mass and isotope simulation of the protonated 2-diethylamino-6-hydroxy-4-methylpyrimidine at m/z $[M + H]^+$ = 182.1288. Metabolite M2 showed a molecular ion peak at m/z $[M + H]^+$ = 182.1285, which is 108 Da lower than M1, consistent with loss of a dimethyl phosphate moiety (O–P(=O) $(OCH_3)_2$) through hydrolysis, forming 2-diethylamino-6-hydroxy-4-methylpyrimidine. Accurate mass data, including elemental composition, theoretical and experimental m/z values, and mass accuracy (ppm), are reported for all compounds and metabolites to support structural assignments.

**Supplementary Table 4. Primers used in this study.**

| Primer name | Sequence 5'-3' | Note |
|---|---|---|
| *Abch2*-NheI-F | CTAGCTAGCTAAAATGTTGATAACCGGCAGTGAGAC | Cloning |
| *Abch2*-XhoI-R | ACCGCTCGAGTTAGCCGCGCTTGAACTTCAG | Cloning |
| *Cyp9k1*-EcoRI-F | ATTGGGAATTCACAACATGTTGGGTACGCTTGC | Cloning |
| *Cyp9k1*-NcoI-R | TCTCCCATGGCTACTTGCGCAACCGGAACC | Cloning |
| *RPS7*-qF | AGAACCAGCAGACCACCATC | qPCR |
| *RPS7*-qR | GCTGCAAACTTCGGCTATTC | qPCR |
| *Abch2*-qF | GCAAGGAGGTGCTCATCAGT | qPCR |
| *Abch2*-qR | TCCCTTCGACGTAAAGTTGGA | qPCR |
| *Cyp6p3*-qF | TGTGATTGACGAAACCCTTCGGAAG | qPCR |
| *Cyp6p3*-qR | ATAGTCCACAGATGGTACGCGGG | qPCR |
| *Cyp9k1*-qF | CAGAAGCGGTCGGTTGACTG | qPCR |
| *Cyp9k1*-qR | AGTTCGTGCGCCATAAATGC | qPCR |
| *Gste2*-qF | CCGGAATTTGTGAAGCTAAACC | qPCR |
| *Gste2*-qR | GCTTGACGGGGTCTTTCGG | qPCR |
| *Coeae6g*-qF | GCTGGAAGGATCGGAAGACTGTCT | qPCR |
| *Coeae6g*-qR | CTCATCCTGGAGCAGATATTCGGGC | qPCR |
